# Coordination of transcriptional and translational regulations in human epithelial cells infected by *Listeria monocytogenes*

**DOI:** 10.1101/775148

**Authors:** Vinko Besic, Fatemeh Habibolahi, Benoît Noël, Sebastian Rupp, Auguste Genovesio, Alice Lebreton

**Affiliations:** Bacterial Infection & RNA Destiny group, Institut de biologie de l’ENS (IBENS), École normale supérieure, CNRS, INSERM, Université PSL, 75005 Paris, France; Computational Biology and Bioinformatics group, Institut de biologie de l’ENS (IBENS), École normale supérieure, CNRS, INSERM, Université PSL, 75005 Paris, France; INRAE, IBENS, 75005 Paris, France

**Author notes:** Equal contribution (Fatemeh Habibolahi, Benoît Noël).

**Keywords:** *Listeria monocytogenes*, Host-pathogen interactions, Translation, 5’-terminal oligopyrimidine motif, Poly(A)-binding proteins

## Abstract

The invasion of mammalian cells by intracellular bacterial pathogens reshuffles their gene expression and functions; however, we lack dynamic insight into the distinct control levels that shape the host response. Here, we have addressed the respective contribution of transcriptional and translational regulations during a timecourse of infection of human intestinal epithelial cells by an epidemic strain of *Listeria monocytogenes*, using transcriptome analysis paralleled with ribosome profiling. Upregulations were dominated by early transcriptional activation of pro-inflammatory genes, whereas translation inhibition appeared as the major driver of downregulations. Instead of a widespread but transient shutoff, translation inhibition affected specifically and durably transcripts encoding components of the translation machinery harbouring a 5’-terminal oligopyrimidine motif. Pre-silencing the most repressed target gene (*PABPC1*) slowed down the intracellular multiplication of *Listeria monocytogenes*, suggesting that the infected host cell can benefit from the repression of genes involved in protein synthesis and thereby better control infection.

## Introduction

Invasion and proliferation of intracellular bacterial pathogens in human cells trigger drastic changes in cell functions, including their gene expression^1^. For instance, the infection of cells by a variety of bacterial invaders has been described to trigger the activation of pro-inflammatory transcription factors, as well as a transient inhibition of host cap-dependent translation^2^. Meanwhile, the survival and multiplication of intracellular bacteria depends upon their capacity to subvert host cell metabolism, functions and antibacterial defences, part of which can be achieved by perturbing host gene expression^3^.

In the past decade, due to the rise of high-resolution transcriptomic approaches, the host transcriptional response to bacterial infections has been extensively explored in a broad range of biological contexts. In contrast, few studies have investigated the perturbation of host translation at an omics scale. Previous reports however support the existence of potent regulations affecting host mRNA translation during bacterial infections. For instance, a growing number of studies have finely characterised miRNA-mediated regulation of specific host transcripts and cellular processes^4,5^. Pathogenic bacteria can also target central host translation mechanisms, and thereby tune —positively or negatively— the production of host defence proteins^6^. The best described example to date is probably that of the intracellular bacterium *Legionella pneumophila* (*Lp*), which has been shown to secrete effectors targeting host translation elongation and thereby stimulate cytokine production ^7^. In line with this, the translatome of *Lp*-infected murine macrophages has pioneered the attempts to discriminate between transcriptional and translational inputs in the fine-tuning of the inducible immune response to infection^8^. Barry *et al*. highlighted that the superinduction of cytokine mRNA transcription enabled infected cells to overcome the general translation elongation blockade imposed by *Lp* effectors, and thus launch a pro-inflammatory response.

*Listeria monocytogenes* (*Lm*) is the foodborne cause of listeriosis, an opportunistic disease of human and cattle that can have severe consequences during pregnancy or in elderly patients. This facultative intracellular bacterium has long been a model for studying all aspects of infection biology, from host-pathogen interactions at the molecular level to *in vivo* and epidemiology studies^9^. How the combination of (*a*) the activity of virulence factors and (*b*) cell autonomous responses contribute to the re-organisation of cell functions has been extensively studied; however, in this model as in others, gene expression has mostly been addressed in terms of transcriptomics, microRNA profiling, activation of pro-inflammatory signalling cascades or chromatin-based regulations^10^. To our knowledge, the effect of *Lm* infection on translation has neither been quantified by mapping the translatome of infected cells, nor by assessing overall changes in protein synthesis rates. As described for other bacterial infections, cap-dependent translation initiation is nonetheless predicted to be transiently impaired during the first hour of infection. Indeed, infection-related stress, and principally membrane pores generated by a secreted perforin, listeriolysin O (LLO), were reported to activate the eIF2α-kinases of the integrated stress response (ISR) pathway and inhibit mTOR signalling, both of which control cap-dependent translation initiation^11–14^. Meanwhile, other *Lm* effectors can restore mTOR signalling; for instance the internalisation protein InlB, by binding the cellular receptor Met, is a potent agonist of growth factors signalling cascades, including mTOR^15^. Signalling pathways coordinating overall cellular translation thus receive positive and negative inputs during infection, which are likely to fluctuate over time. On top of these, specific translational regulations may control the expression of defined subsets of genes, downstream of transcriptional regulation. Ultimately, the possible consequences of modulating host translation on *Lm* infection outcome have not been explored.

In the present study, we aimed at clarifying the respective contribution of transcriptional and translational regulations on the reshaping of host gene expression of a human epithelial cell line, over a 10-h time course of infection with an epidemic isolate of *Lm*. Using ribosome profiling^16^, we mapped with high resolution the host translatome during infection, compared it with transcriptome data, and grouped genes that were under transcriptional and/or translational control according to their regulation profiles with regards to time. Our results revealed a dominant pattern, where the rapid induction of gene expression was mainly driven by transcriptional regulation and affected inflammation-related genes, whereas most repressive events were translational, and affected genes encoding components of the translation machinery. The most repressed gene was *PABPC1*, encoding the host cytoplasmic poly(A)-binding protein. Interestingly, preventing *PABPC1* expression by using siRNA-mediated silencing dampened the replication of *Lm*, suggesting that limiting the expression of genes involved in the translational machinery could be part of the cellular responses that help cope with the severity of infection.

## Results

### *Listeria* infection does not significantly impair the translation capacity of epithelial cells during the first ten hours of infection

In contrast to what has been shown for other pathogenic bacterial species^2^, whether infection by *Lm* affects the overall translation activity of host cells was unknown. To assess this, we quantified the ability of a human epithelial cell line from a colon adenocarcinoma (LoVo) to incorporate the methionine analogue homopropargylglycine (HPG) into newly synthesised proteins, over a ten-hour time course of infection by a strain of *Lm* from an epidemic isolate, LL195^17,18^ (*Figure 1A*). Infection conditions with this hypervirulent strain were optimised in order to maximize the proportion of infected and viable cells over the period considered. When using a multiplicity of infection (MOI) of 30, bacterial intracellular growth was exponential until 10 h post-infection (p.i.), and the loss of viability of host cells was minimal (*Figure S1A*). More than 40% of cells were infected as soon as 2 h p.i. and then, due to cell-to-cell spread of bacteria, the infection expanded so that more than 90% of cells were infected at 5 h p.i (*Figure S1B*), which can be considered sufficiently homogeneous for analysing translational effects on cell populations. To avoid possible side effects on host translation due to the change of medium composition in nutrients and growth factors, all experiments were performed using “conditioned medium”, in which cells had been grown for one day before it was pooled and saved. The doubling time of LoVo cells in conditioned medium remained similar to usual culture conditions when the medium is changed 2-3 times a week (34 h), and entry rates when using conditioned medium with a MOI of 30 were similar to that obtained in medium that does not contain serum and a MOI of 20.

**Figure 1.**
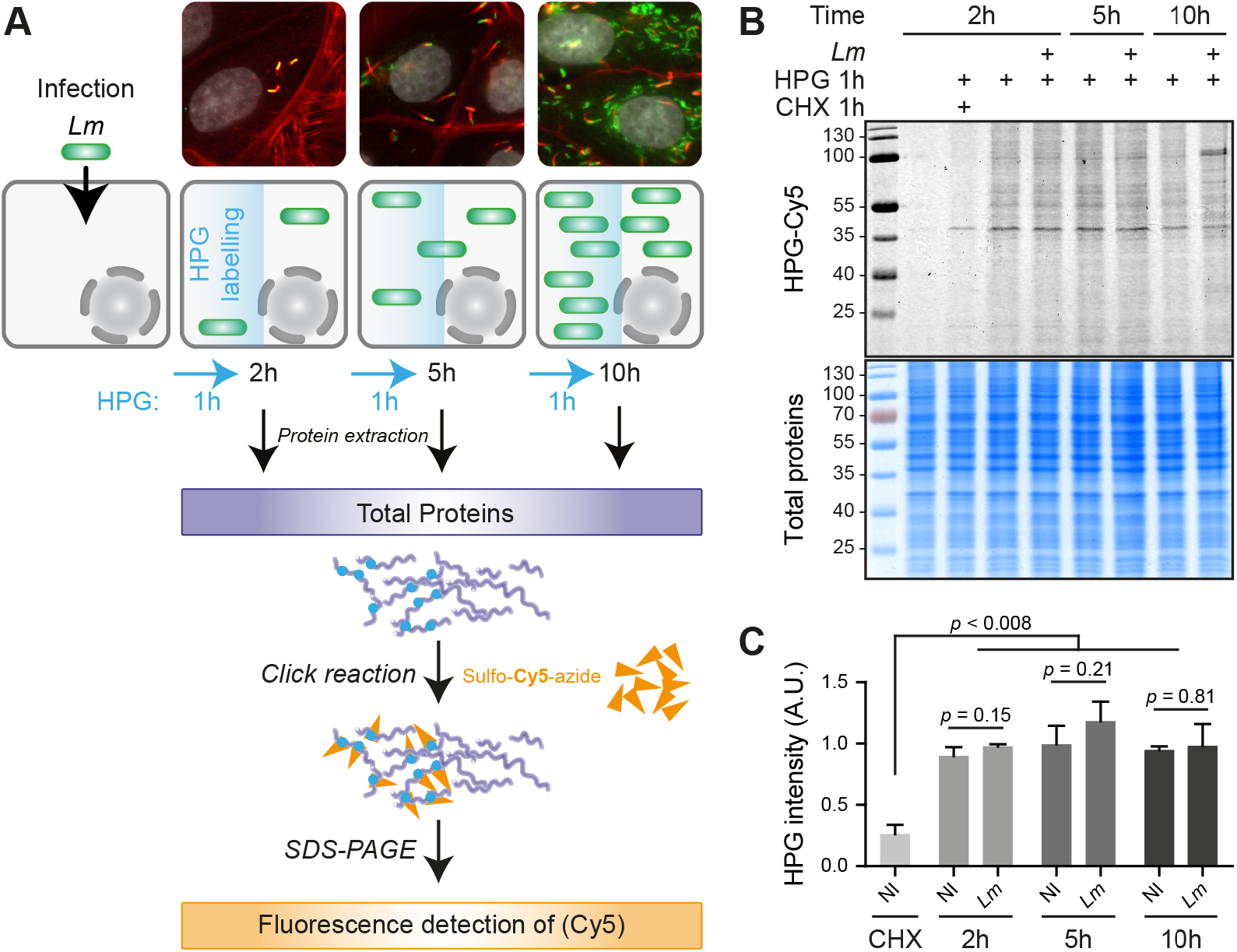
*Lm* infection has a low impact on total translation activity in LoVo epithelial cells. (A) Principle of the metabolic labelling of newly synthesized proteins with homopropargylglycine (HPG) over an infection time course. LoVo cells, infected or not for 2, 5 or 10 h with *Lm* LL195 constitutively expressing eGFP, were treated with HPG for 1 h prior to recovery. Cell infection was monitored by immunofluorescence staining on coverslips. DAPI staining of cell nuclei is displayed in white, F-actin staining by fluorescently-labelled phalloidin is in red, and eGFP-expressing bacteria are in green. After cell lysis, HPG residues that had been incorporated into newly-synthesised proteins were conjugated with sulfo-Cy5-azide by copper-catalysed alkyne-azide cycloaddition. (B) In-gel fluorescence detection of HPG incorporation into newly synthesised proteins. Following cycloaddition, protein samples were separated by SDS-PAGE, and Cy5 fluorescence was recorded (*top panel*), before the gel was stained with colloidal Coomassie as a loading control (*Bottom panel*). (C) Quantification of HPG incorporation. The integrated density of Cy5 fluorescence was measured for each lane and normalized to the corresponding integrated density of Coomassie staining. NI, non-infected. Data are average and standard deviation from independent experiments; n = 3 for CHX, 2 and 5 h p.i.; n = 2 for 10 h p.i. *p*-values were calculated by two-tailed T-tests.

To evaluate the efficiency of protein synthesis, HPG was added to uninfected or infected cell cultures, one hour prior to each recovery time point, then the labelling of newly-synthesized proteins was revealed by cycloaddition of HPG residues with sulfo-Cy5-azide, followed by sodium dodecyl sulfate-polyacrylamide gel electrophoresis (SDS-PAGE), in-gel fluorescent detection (*Figure 1B*) and quantification (*Figure 1C*). Compared to non-infected cells, no major difference was observed in the overall intensity of HPG incorporation in infected cells at 2, 5 or 10 h p.i compared to non-infected conditions (NI). Whereas total amounts of newly synthesized proteins appeared grossly unchanged, at 10 h p.i., the pattern of labelled proteins started differing, essentially due to the accumulation of an abundant, newly-synthesised protein of ~100 kDa. As expected, treatment of one of the samples with cycloheximide (an inhibitor of translation elongation) concomitantly with HPG addition blocked HPG incorporation, arguing that the detected signals were representative of the cellular activity of protein synthesis.

*Lm* LL195 thus does not seem to impose a noticeable translational shutoff on its host, but tends on the contrary to maintain a level of protein synthesis comparable to that of uninfected cells. Based on this result, we considered that a translatome analysis of the infected cells could be undertaken, without running the risk that normalizing sequencing data to library size would mask overall changes of translation rates.

### Early host gene expression response is dominated by transcriptional activation, while repression events are mainly translational

To clarify whether specific host transcripts were the object of translational regulation during infection, we assessed mRNA expression and translation using high throughput Illumina mRNA sequencing (hereafter, RNA-seq) and ribosome profiling (hereafter, Ribo-seq) from biological samples in triplicates that were either non-infected, or recovered at 2, 5 or 10 h p.i. As expected from this technique, RNA-seq generally had a higher amount of uniquely mapped reads (24 to 32 million) than the Ribo-seq (*Figure S2A*). Almost all Ribo-seq samples had nearly 10 million reads or more, except replicate #2 at the 10 h time-point, which not only had less than three million reads, but also had a smaller proportion of reads mapping to coding sequences (CDS). Consequently, this sample was considered of poor quality and subsequently removed from downstream analysis. Sequenced ribosomal footprints (RFPs) displayed the expected length profile, which peaked at 29 nucleotides (*Figure S2B*), typical of high quality Ribo-seq data sets. As expected from Ribo-seq data, RFPs also mapped predominantly to the coding sequences (CDSs) and 5’-untranslating regions (UTRs), with little mapping to the 3’-UTRs (*Figure S2C*). Moreover, mapped RFPs displayed the three-nucleotide codon periodicity characteristic of translating ribosomes (*Figures S2D-E*). Infection appeared to have no drastic effect on average translation elongation profiles (*Figure S2D*), thus confirming the conclusions drawn from HPG labelling experiments, that overall translation activity is barely perturbed by a 10-h *Lm* infection in epithelial cells.

The number of uniquely-mapped reads, then Reads Per Kilobase of transcript, per Million mapped reads (**RPKM**) values for each gene was computed for each RNA-seq or Ribo-seq sample, and Pearson correlation coefficients of RPKM values were calculated between each pair (*Figure S3*). Apart from Ribo-seq replicate #2 at 10 h p.i. that we discarded, maximum correlation coefficients (above 0.97) were observed between all RNA-seq samples on the one hand, and all Ribo-seq samples on the other hand. At any given time-point, correlation coefficients between replicates in RNA-seq or Ribo-seq were between 0.98 and 1.

We computed the fold changes in abundance of transcripts between each time point, either in the RNA-seq or in the Ribo-seq datasets. The Spearman correlation coefficients between log_2_ fold changes (FC) among differentially regulated genes (DRGs) in the RFP *vs* RNA values were 0.906, 0.771 and 0.759 at [2 *vs* 0], [5 *vs* 2], and [10 *vs* 5] h p.i respectively. The corresponding scatter plots are represented as *Figure 2A-C* and source data is provided in *Table S1*.

**Figure 2.**
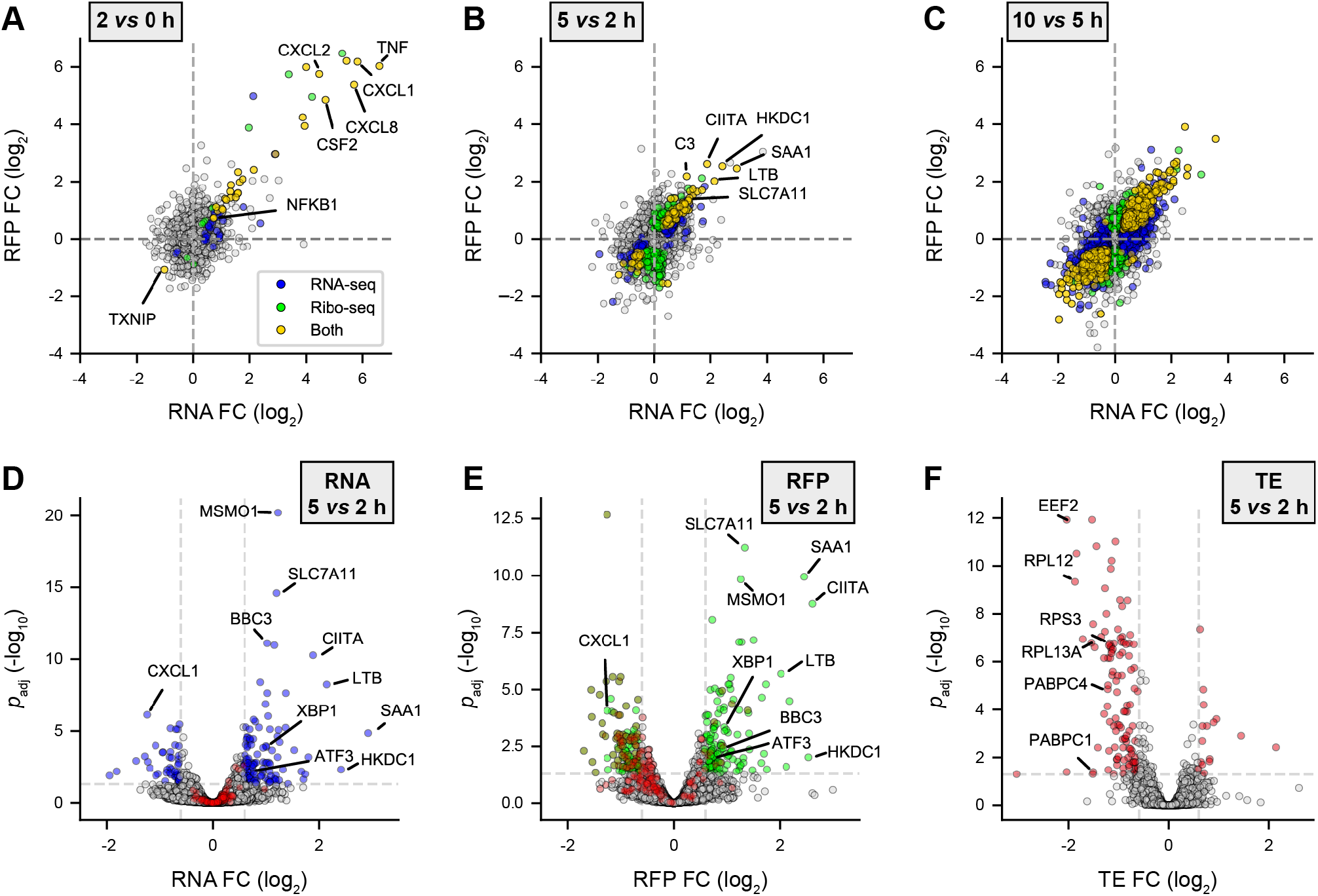
Transcriptional up- & translational down-regulations dominate gene expression response to *Lm* in the first hours of infection. LoVo intestinal epithelial cells were infected for 2 to 10 h. Cell lysates were processed for total cytoplasmic RNA-seq and ribosome profiling (Ribo-seq). (A-C) Scatter plots of changes in normalized RFP (*Y axis*) *vs* RNA (*X axis*) levels along the course of infection, when comparing (A) 2 h *vs* non-infected, (B) 5 *vs* 2 h, or (C) 10 *vs* 5 h. (D-E) Volcano plots highlighting genes being significantly up- (*right*) or down- (*left*) regulated in (D) RNA-seq, (E) Ribo-seq, or (F) translation efficiency (TE) at 5 h p.i. compared to 2 h p.i. Data points coloured in blue, green or red represent genes with *p*_adj_ below 0.05 (above dashed grey horizontal line; −log_10_ *p*_adj_ = 1.3) and a FC below or above 1.5 (vertical dashed grey lines; log_2_ FC = ± 0.58). Genes for which *p*_adj_ < 0.5 in TE are highlighted in red in D and E. (A-E) Data from three independent replicates (except for RFPs at 10 h). FC, fold change; *p*_adj_, adjusted *p*-value [DESeq false discovery rate (FDR)].

Comparing the 2-h infected time-point with the non-infected controls, DESeq2 differential expression analysis detected 68 DRGs in the RNA-seq and/or Ribo-seq datasets (adjusted *p* value, *p*_adj_ < 0.05) (*Figure 2A*). Most of these genes clustered next to the diagonal in the upper right quadrant of a Ribo-seq *vs* RNA-seq scatter plot, highlighting that the dominant feature of gene expression changes during the first two hours of infection was the transcriptional induction of a subset of host transcripts, which then underwent translation without further translational control. Indeed, out of 29 genes displaying a similar differential regulation at the levels of RNA and RFP, all but one were upregulated, with fold changes ranging from 1.7 to 70. *TXNIP*, an inhibitor of anti-oxidative pathways, was the only gene to be statistically significantly repressed (FC: −2, *p*_adj_ = 4.2 × 10^-7^ for RNAs, and 2.3 × 10^-3^ for RFPs).

Between 2 and 5 h p.i., a total of 621 genes showed differential regulation, in either or both of the datasets (*Figure 2B*). 112 (18%) of these DRGs displayed a similar trend in both the RFP and RNA patterns, indicating that their regulation was driven by changes in their RNA synthesis or decay rates. 319 transcripts (58% of DRGs) appeared as only significantly regulated in the translatome dataset. The overlap between DRGs that were significantly deregulated (*p*_adj_ < 0.05) and displayed fold-changes above 1.5 in the Ribo-seq versus RNA-seq datasets is displayed as Venn diagrams in *Figures S4A* (down-regulated genes) and *S4B* (up-regulated genes). These intersections reveal a higher overlap between RNA and RFP datasets for up-regulations (59 genes, 28.38% of DRGs) than for down-regulations (14 genes, 7.87% of DRGs).

Between 5 and 10 h p.i., 4,537 genes were further deregulated, among which 1,078 genes (24%) were similarly regulated in both RNA and RFP datasets (*Figure 2C*). All types of differential regulations (positive or negative, affecting the transcriptome or the translatome) were identified, arguing that over time various regulatory mechanisms cooperate to best adapt the expression of each host gene to changing conditions. Note that, due to the loss of one of the translatome samples at the 10 h time points, the proportion of transcripts that qualified as significantly regulated for RFPs in this dataset is likely underestimated.

In order to analyse variations in the translation of each transcript independently of transcript abundance, we used Riborex to compute changes in translation efficiency (TE) during the course of *Lm* infection (*Figure 2D-F, Figure S5, Table S1*). We analysed more extensively changes in TE that occurred between 2 and 5 h p.i. as being the most prominent, and addressed whether they were driven by transcriptional and/or translational regulations. *Figures 2D-F* displays the volcano plots of changes in RNA counts, RFP counts or TE values, between 2 and 5 h p.i. The genes for which changes in TE were significant (*p*_adj_ < 0.05, in red) were little affected by variations in the RNA-seq dataset (*Figure 2D*). In contrast, the majority of DRGs for TE were grouped on the left side of the Ribo-seq volcano plot (*Figure 2E*), arguing of translational repression. No gene that changed at the RNA level did not also change at the RFP level among the statistically-significant genes for TE variation. Regarding RNA-seq data, 121 out of 160 (75.6%) of the significant DRGs with FC > 1.5 were upregulated (*Figure 2D, right, Figure S4B*). 48.8% of these were also positively regulated in the RFP data (*Figure 2E*, *right, Figure S4B*), corresponding to the genes that grouped next to the diagonal in the upper right quadrant on *Figure 2B*, and for which upregulation of protein synthesis correlated with increased transcript abundance without change in TE. In contrast, only 14 (9.8%) out of the 143 down-regulated genes in the Ribo-seq dataset were also repressed in the RNA-seq dataset (*Figure 2D-E*, *left, Figure S4A*), indicative of the predominance of repressive events at the translational level over transcriptome level. These repressive translational events are more clearly illustrated by TE data on *Figure 2F*, where the vast majority of genes that were affected in TE grouped in the left part of the volcano plot.

Altogether, this analysis confirmed that, while the positive regulation of host gene expression in the first five hours of infection was mainly driven by changes affecting the transcriptome, a subset of genes was affected by translational repression, which was sharply detected between 2 and 5 h p.i.

### Transcriptional induction and translational repression affect functionally distinct biological processes

We then sought to investigate whether genes that were subject to similar changes in their gene expression also shared functions that might prove relevant to the infectious process. To this end, we performed overrepresentation analysis (ORA) of gene ontology (GO) biological processes among DRGs which were either up- or down-regulated in RNA-seq, Ribo-seq or TE at each time point of the infection, compared to the noninfected condition (*Figure 3A, Table S2*). The early transcriptional activation highlighted in *Figure 2* led to a pronounced induction of genes associated with pro-inflammatory and type I interferon responses to bacterial invasion (*Figure 3A*, first seven lines). As expected, this transcriptional activation was directly mirrored by an increased in Ribo-seq data, and persisted through to the 5-h time point, when activation of genes related to autophagy, apoptosis and ER stress additionally occurred. At 10 h p.i., the up-or down-regulation of other pathways started emerging, including protein catabolic response, chromatin silencing and mitochondrial metabolism.

**Figure 3.**
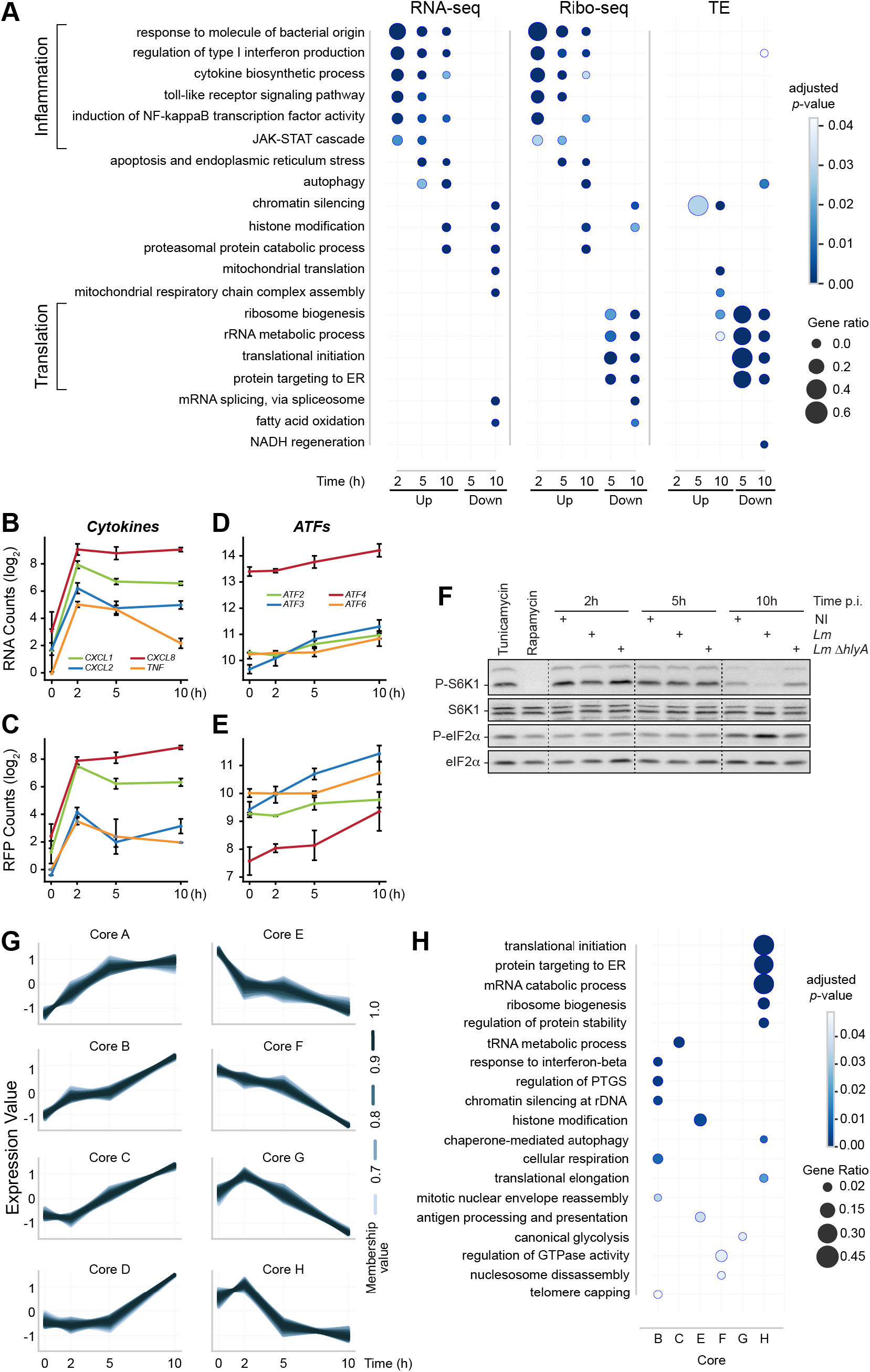
Early transcriptional regulation of inflammatory response precedes a translational repression of the translational equipment. (A) Over-representation analysis (ORA) of GO Biological Process terms for up- or down-regulated genes in RNA-seq, Ribo-seq or TE over all time points. For each time point, DRGs were selected by comparison to the non-infected condition. (B-F) Induction of inflammation and of the integrated stress response by *Lm* infection of LoVo cells. The variation of RNA (B, D) and RFP (C, E) levels was quantified for selected NF-kB transcriptional target genes related to inflammation (B, C) or genes involved in the integrated stress response (D, E) during infection. Data represent DESeq normalized read counts from three independent experiments and error bars indicate standard deviation. (**F**) The phosphorylation status of the mTOR substrate S6K1, and of the target of ISR eIF2 kinases eiF2α were assessed by immunoblotting in cells infected for 2, 5 or 10 h by wild-type *Lm*, or by a *Lm* strain where the *hlyA* gene encoding LLO had been deleted (Δ*hlyA*). Treatments by tunicamycin and rapamycin were used, respectively, as an inducer of the ISR through endoplasmic reticulum stress, and as a repressor of mTOR activity. **(G)** Transcripts sharing similar TE profiles over time were clustered by fuzzy clustering. For each cluster, only genes having more than 70% membership are displayed. (**H**) Functional categories were assigned to each cluster by ORA of GO Biological Process terms.

The rapid transcriptional induction of pro-inflammatory cytokine and type I interferon genes in response to *Lm* infection has been largely documented^19^. The sensing of pathogen-associated molecular patterns (PAMPs) by cell sensors is known to activate the transcription factor role of NF-κB. We analysed the enrichment for transcription factor binding sites (TFBS) in the promoter regions of the genes that were significantly transcriptionally induced at 2 h p.i (*Table S3*). Unsurprisingly, binding sites for NF-kB subunits NFKB1/NFKB2/RELA dominated, with normalized enrichment scores (NES) above 9 confirming that the early induction of NF-kB-dependent signalling during *Lm* infection of epithelial cells drives the inflammatory response. The effects of this early transcriptional activation (followed by a rapid downregulation) on the RNA and RFP profiles of a subset of cytokine genes is illustrated in *Figures 3B-C*. At later time points, the weight of NF-kB-dependent transcription declined and the action of additional transcription actors appeared. Between 5 and 10 h p.i., the most enriched motifs upstream of transcriptionally induced genes were binding sites for the stress-responsive transcriptional factors ATF2, ATF3 and ATF6 (NES above 4, *Table S3*), highlighting contribution of the ISR to the host transcription as infection proceeded. In line with this, the expression of these three transcription factors rose over time (*Figures 3D-E*). By immunoblotting, we confirmed that eIF2α phosphorylation increased in the latest time-points of infection, suggesting that the ISR eIF2 kinases had been activated (*Figure 3F*). This activation was not detected when cells were infected with a strain that did not produce LLO, confirming previous reports that the major pore-forming toxin of *Lm* is a critical determinant of host cell stress responses^13^. Altogether, our data confirm an early transcriptional induction of inflammatory pathways in response to infection, which we find to be followed by the activation of the ISR.

Downregulation of functional pathways only emerged between 2 and 5 h p.i. and was largely dominated by translational repression, noticeable in the comparison of Ribo-seq data, and even more obviously when analysing changes in TE. Strikingly, the majority of the transcripts affected by translational repression encoded proteins involved in translation itself (*Figure 3A*, lines 14-17). At 10 h post infection, transcripts encoding translation components were further downregulated translationally, while there was also a modest decrease in the TE of a few genes regulating the type I interferon response, autophagy and NADH regeneration. The decrease in TE of genes involved in NADH regeneration was mostly explained by a decrease in RFPs suggesting repressive translational mechanisms, whereas the apparent decreased TE of genes involved in autophagy and in the regulation of the type I interferon response reflected a higher increase in their transcript abundance than in their RFPs, perhaps suggesting a buffering effect on their translation.

### Genes that are translationally co-regulated over time group into functionally-related clusters

To investigate the effect of time on host translation during *Lm* infection, we conducted fuzzy clustering on genes displaying a differential TE in the Riborex analysis, using a cut-off of 0.1 for the adjusted *p* value. Eight core clusters were generated, representing the major temporal patterns of TE changes during *Lm* infection (*Figure 3G, Table S4*). Four of these cores (A to D) contained 393 genes that showed an overall increase in TE over the ten-hour time course, while the remaining four cores (E to H) contained 525 genes that displayed an overall decreased TE. In order to assess the functional relevance of these patterns of TE changes, we conducted ORA analysis of GO biological processes on each one of the cluster cores. Six of the cores were statistically enriched for specific GO biological processes (*Figure 3H, Table S4*), which are detailed hereafter.

Transcripts belonging to core B displayed a steady increase in TE throughout the infection and encoded factors involved in distinct biological processes. These included regulation of silencing, either chromatin-based or post-transcriptional, but also genes involved in host response to pathogens, such as type I interferon response, or cellular respiration. Core C was marked by an increase in TE starting after 2 h p.i., and was enriched for genes involved in non-coding RNA metabolism, and predominantly “tRNA metabolic processes”.

Core E, in which transcripts were affected by a steady decrease in TE throughout the infection, was enriched for “histone modification” processes and to a lesser extent, “antigen processing and presentation of exogenous antigen” processes, which mostly represent proteins involved in cargo targeting to vesicles and their processing. In Core F, transcripts were affected by a mild decrease in TE that intensified between 5 and 10 h p.i. It was moderately enriched for processes representing “nucleosome disassembly” and “regulation of GTPase activity”. All the genes belonging to the “nucleosome disassembly” process encoded proteins participating in transcriptional regulation via chromatin remodelling. Among them, SMARCA4 had been previously reported to exert a repressive role on E-cadherin transcription^20^. Genes belonging to the “regulation of GTPase activity” process mainly encoded regulators of actin dynamics or vesicle formation and processing. Altogether cores E and F, both of which present a temporal decrease in TE, are broadly enriched for genes involved in chromatin-based regulation, vesicle formation and processing.

Cores G and H were characterized by a slight increase in TE at 2 h p.i. followed by a prominent decrease between 2 and 5 h p.i., which was further amplified until 10 h in core G whereas it plateaued after 5 h in core H. In both of these clusters, changes in TE were largely due to a strong decrease in the abundance of RFPs (rather than by an increase in RNA-seq), suggesting these genes were actually translationally repressed rather than buffered by a lack of translation. Core G was moderately enriched for genes related to the “regulation of canonical glycolysis”. In contrast, core H contained essentially genes encoding factors involved in ribosome biogenesis and protein synthesis; indeed, 80% of them encoded ribosomal proteins, translation initiation or elongation factors. A few additional genes were associated with related biological processes, such as “mRNA catabolism”, “protein stability”, and “autophagy”.

In terms of number of co-regulated genes, the enrichment of core H with factors required for host translation was the most striking feature of the fuzzy clustering and ORA analysis. We then asked whether specific features within these functionally-related transcripts could determine their co-regulation.

### 5’-terminal oligopyrimidine motif-containing mRNAs are co-repressed translationally during *Listeria* infection

The length and sequence of transcript 5’-UTRs are known to play important roles in the regulation of translation initiation^21^. Here, no specific trend linking the length of transcript 5’-UTRs and overall Log2 FC in TE over time was observed (*Figure S6*) even though we cannot formally exclude their contribution to the observed translational regulations. Several *cis*-acting motifs located within the 5’-UTRs of mRNAs have previously been described for their ability to modulate eukaryotic translation initiation in response to cellular stresses or infection. Among them, four classes of motifs can have a significant influence on the efficiency of recruitment of the translation pre-initiation complex (PIC) or of initiation at the appropriate AUG, namely: translation initiator of short 5’UTRs (TISU), 5’-terminal oligopyrimidine motifs (5’-TOP), internal ribosome entry site (IRES) structures, or upstream open reading frames (uORF)^22^. To assess whether the presence of these motifs could dictate the co-regulation of the transcripts that were translationally repressed during infection, we tested whether the repressed gene set displayed a statistically significant enrichment for any of these motifs using ROAST. Out of the 1,003 genes that had differential TE across any one condition as calculated by Riborex, 82 were only translationally regulated and had more than twofold change in RFP levels across the ten-hour time-course, while their transcript abundance remained stable. We found that there was a significant enrichment of TOP genes at 10 h p.i. (ROAST *p*_adj_ = 0.021) in this set, whereas none of the other motifs were statistically enriched.

To further illustrate this enrichment, we displayed the distribution of changes in TEs during infection for each list of experimentally validated TOP- (n = 83), uORF- (n = 76) or IRES- (n = 25) containing transcripts, or for the 133 transcripts containing a TISU motif with no mismatch (*Figures 4A-D, Table S5*). After a modest increase in TE for the bulk of TOP-containing transcripts at 2 h p.i., they consistently displayed a decrease in TE at 5 h p.i., which exacerbated at 10 h p.i. (*Figure 4A*). In contrast, the distributions of uORF-, IRES- or TISU-containing transcripts remained centred on 0 (*Figures 4B-C*). The individual profiles for each one of the transcripts belonging to each category revealed individual variability within the general trends (*Figures 4E-H, Table S5*), and emphasised a high similarity in profiles between TOP-containing transcripts (*Figure 4E*) and core cluster H encoding genes involved in host translation (*Figures 3G-H*). Altogether, our data indicate that the major common feature of the transcripts encoding translation-related proteins that were translationally repressed during infection was the presence of a TOP motif in their 5’-UTR.

**Figure 4.**
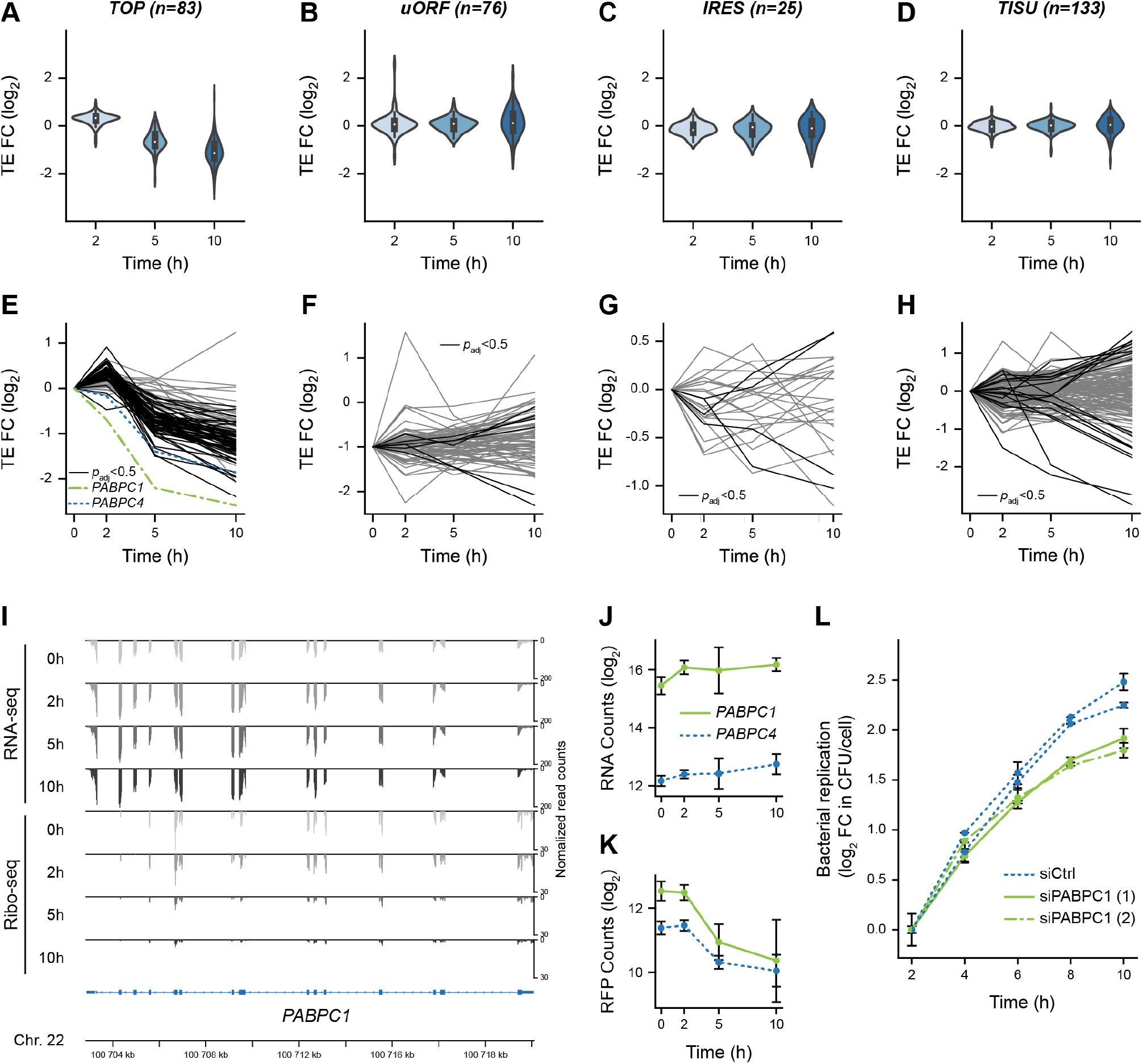
Translational repression of 5’-terminal oligopyrimidine-containing transcripts, including *PABPC1*, during *Lm* infection. (A-D) Violin plots representing fold changes in TE across time of transcripts that have been experimentally verified to contain (A) functional TOP, (B) uORF, or (C) IRES motifs, or (D) predicted to contain a TISU in their 5’-UTR regions. (E-H) Translation efficiency profiles of individual transcripts containing either a (E) TOP, (F) uORF, (G) IRES or (H) TISU motifs. Transcripts for which the adjusted *p*-value from Riborex analysis was below 0.5 are displayed in black line (except *PABPC1* and *PABPC4*, in dotted green and blue lines, respectively), while transcripts for which TE changes were not significant are displayed in grey. (I) Profiles of RNA-seq (*top*) and Ribo-seq (*bottom*) reads aligned at the *PABPC1* locus. Average values of read counts per genomic position from three independent experiments, normalized for library size, are represented for each time-point. (J-K) Quantification of the variation of *PABPC1* and *PABPC4* RNA (J) and RFP (K) levels during infection. Data represent DESeq normalized counts from three independent experiments and error bars indicate standard deviation. (L) Silencing of *PABPC1* reduces *Lm* intracellular replication rate. LoVo cells were transfected with siRNA against *PABPC1* mRNA or a scrambled siRNA (siCtrl) for 48 h before infection. Two independent experiments were carried out using distinct siRNA against PABPC1 (siPABPC1 (1) and (2)). Bacterial entry and replication were assessed by gentamicin protection assay followed by serial dilution plating of infected cell lysates on agar plates. In order to focus on intracellular multiplication rather than differences in entry, the log_2_ ratio of colony forming units (CFU) counts per cell at each time points relative to time 2 h post-infection were plotted. Values are averages and standard deviation from three infected wells per condition, each counted thrice.

Because TOP-containing transcripts have been previously shown to be translationally repressed in conditions were the activity of the mTOR kinase was impaired^23^, we probed for the phosphorylation status of the mTOR substrate S6 kinase 1 a (S6K1) on serine 51 by immunoblotting (*Figure 3F*). Whereas rapamycin treatment of LoVo cells for one hour was sufficient to abrogate S6K1 phosphorylation, we only observed a complete dephosphorylation of this substrate after 10 h of *Lm* infection, arguing that loss of mTOR activity alone could not account for the translational repression of TOP-containing transcripts that we observed as soon as 5 h p.i.

Downstream of mTOR signalling, the main actor of the translational regulation of TOP genes has recently been shown to be LARP1, which would either bind the TOP motif and repress translation when it is unphosphorylated, or allow translation when it is phosphorylated^24–26^. To assess whether the impaired translation of TOP genes could be controlled by LARP1 dephosphorylation in infected LoVo cells, we assessed its phosphorylation status by immunoprecipitating LARP1, and then probing with a total anti phosphoserine/threonine antibody (*Figure S7*). Whereas LARP1 was efficiently pulled-down, in our hands it did not appear to be phosphorylated in non-infected LoVo cells, suggesting that the translation of TOP genes at 0 h p.i. was independent of LARP1 phosphorylation. We thus assume that another layer of regulation than LARP1 dephosphorylation might be at play for the repression of TOP genes observed in infected LoVo cells.

### *PABPC1* is translationally repressed during *Listeria* infection of epithelial cells

Interestingly, the only two TOP-containing transcripts that displayed a decrease in TE as soon as 2 h p.i. were *PABCP1* and *PABPC4*, with *PABPC1* being the most heavily repressed of all TOP-containing transcripts (*Figure 4E*). *PABPC1* and *PABPC4* encode cytoplasmic poly(A)-binding proteins (PABP), and are the only two members of this family to be expressed in LoVo cells. Both transcripts underwent a potent translational inhibition during *Lm* infection, which deepened as the infection proceeded in spite of a slight increase in their transcript abundance over time (*Figures 4E and 4I-K*). Because *PABPC1* was the most expressed of the two paralogues and displayed the most striking repression, we focussed on the regulation of the expression of this gene during *Lm* infection. Using fluorescence *in-situ* hybridization (FISH) against *PABPC1* mRNA, we confirmed that its abundance did not decline in infected cells, even though a noticeable heterogeneity between individual cells could be noticed at all time points (*Figure S8*). The cytoplasmic location of *PABPC1* transcripts was generally diffuse, and a small proportion of the signal co-localised with the P-body marker DDX6. No remarkable change in the proportion of *PABPC1* mRNA localising to P-bodies was observed throughout the infection time-course, arguing that sequestration of the transcript within these compartments could likely not account for the intensity of the translational repression detected in our analysis.

### De-regulation of *PABPC1* expression impacts on *Listeria* intracellular replication

The potent control exerted on *PABPC* mRNA translation prompted us to scrutinize if cytoplasmic *PABPC1* translation or abundance was having an impact on infection progression. Data from a previously published siRNA screen for host factors involved in *Lm* infection of HeLa cells indicated that silencing of *PABPC3* (which is not expressed in LoVo cells), and to a lesser extent *PABPC1*, reduced intracellular bacterial loads at 5 h p.i.^27^. We hypothesised that reducing the synthesis of PABPCs during infection might facilitate the ability of infected cells to control bacterial intracellular multiplication. To test this hypothesis, we transfected LoVo cells with siRNAs against *PABPC1* (two distinct siRNAs were used), or with a control scrambled siRNA, 48 h before infecting with *Lm*, and then monitored the intracellular replication of bacteria in cells (*Figure 4L*). Intracellular bacteria were recovered by cell lysis and plated on BHI-agar plates at several times post-infection, then colony forming units (CFU) counts were normalized to values measured at 2 h p.i. in order to analyse only the intracellular multiplication of bacteria rather than possible variations in entry into cells. In parallel, the efficiency of *PABPC1* silencing in transfected cells was assessed by immunoblotting (*Figure S9A*). When *PABPC1* was silenced in LoVo cells, the intracellular replication rate was significantly (though modestly) reduced, compared to cells transfected with a control siRNA (*Figure 4L*). These observations suggest that repressing the expression of cytoplasmic PABPs could participate in cellular control of bacterial proliferation.

## Discussion

Regulation of gene expression allows organisms to respond to changes in their environment. The intensity and kinetics of the response is strongly influenced by the nature of the regulatory mechanisms involved, affecting various levels on the path from DNA to end-products. In the present study, we aimed at clarifying the respective contribution of transcriptional and translational regulations on the reshaping of host gene expression of a human epithelial cell line, over a 10-h time course of infection with an epidemic isolate of *Lm*. Metabolic labelling with homopropargylglycine revealed that *Lm* infection did not drastically impair the overall translation capacity of infected epithelial cells. By comparing translatome with transcriptome data, we then identified genes that were under transcriptional and/or translational control, and grouped them according to their regulation profiles with regards to time. Our results revealed a dominant pattern, where the rapid induction of gene expression was mainly driven by transcriptional regulation, whereas most repressive events were translational. Over-representation analysis of gene ontologies also highlighted a frequent co-regulation of genes encoding proteins involved in related biological processes. Typically, whereas inflammation was transcriptionally induced, most genes encoding components of the translation machinery were translationally repressed, likely due to a strong repression of the translation of mRNAs harbouring a 5’-terminal oligopyrimidine (TOP) motif. The most repressed gene was *PABPC1*, encoding the main host cytoplasmic PABP. Interestingly, further repressing *PABPC1* expression using siRNA-mediated silencing dampened the replication of *Lm*, suggesting that limiting the expression of the translational machinery could be part of the cellular responses that helps the cell cope with the severity of infection.

### Contribution of infection-induced stress responses to gene expression regulation

In addition to the expected early activation of NF-kB and its pro-inflammatory effect, a large part of the regulations we observed could be explained by the inhibition of mTOR signalling and the activation of the ISR. Both were previously described in response to *Lm* infection, and were mainly dependent on the pore-forming activity of LLO^12–14^. Treatment of RPE1 cells by LLO was also shown to trigger a transient phosphorylation of eIF2α—a hallmark of ISR—, as well as a transient arrest in total protein synthesis^11^. However, our results appear to differ significantly from the existing literature in terms of kinetics and intensity. First, no transient arrest in overall protein synthesis occurred during the course of our 10-h infection of LoVo cells (*Figure 1*), suggesting that the drastic effects on protein synthesis observed when cells were treated with an elevated dose of LLO (0.5 or 1 μg/ml, *i.e*. 9 to 18 nM) was not representative of the real exposure of cell membranes to the toxin when it was secreted by invading bacteria. Second, translational and transcriptional effects that could be attributed to the induction of the ISR appeared gradually over time, and became noticeable in transcriptome data only after 10 h of infection, matching an increase of eIF2α phosphorylation at this later time-point (*Figure 3F*). The translation of *ATF4*, which is known to be induced upon phosphorylation of eIF2α due to the presence of a series of uORFs in its 5’-UTR, only modestly increased during infection. The transcription and translation of *ATF3* increased gradually over time since the beginning of infection, while the induction of *ATF6* only occurred between 5 and 10 h p.i. In line with this, the transcriptional targets of ATF3 and 6 were upregulated at 10 h p.i. These observations suggest that in the context of our study, ISR induction by *Lm* infection is gradual and long-lasting, in agreement with what has been previously shown for the phosphorylation of eIF2α due to activation of the unfolded protein response by *Lm*^13^. This conclusion contrasts however with other reports that eIF2α phosphorylation was early and transient^12,14^. Part of this discrepancy may have arisen from noticeable differences in our experimental setup compared to that used by Tattoli *et al*. For instance, we used a lower MOI (30 rather than 100) and lower centrifugation speed (1.5 minutes at 200 × *g* rather than 1 minutes at 2,000 × *g*) than Tattoli *et al*. We have also used conditioned medium throughout our experiments to avoid any possible effects on the sensing of amino-acid starvation when replacing media. We hypothesize that our milder conditions of cell culture and infection could be responsible for delaying and lengthening the effects of the ISR activation reported by some of the other groups.

The metabolism and equipment of the LoVo cell line we have used is also of relevance to its response. This colon adenocarcinoma cell line, which is an acknowledged model for the infection of intestinal cells by *Lm*^28,29^, grows relatively slowly (doubling time: 34 h) and displays reduced total translation activity compared with the HeLa cells or mouse embryonic fibroblasts (MEFs) that were used by Tattoli *et al*.; Pillich *et al*. used P388D1 mouse monocytes, and did not reproduce their observation in HeLa cells; Shrestha *et al*. used RAW 264.7 murine macrophages and MEFs. Another important difference between our experimental conditions and that of others was the use of an epidemiological isolate of *Lm*, the biology of and host response to which have scarcely been addressed to date. The selected strain LL195 (serotype 4b) belongs to the clonal complex 1 (CC1) within lineage I of *Lm*^18^, which is more representative of clinical cases of listeriosis than usual lineage II laboratory strains such as EGD-e (CC9, serotype 1/2a) that was used by Pillich *et al*., or 10403S (CC7, serotype 1/2a) that was used by Tattoli *et al*. The haemolytic titre we measured for LL195 was in the same range as that of EGD-e, arguing against a lack of LLO activity in this strain (*Figure S10*). However, we cannot exclude that a different repertoire of virulence factors expressed by this strain might impact the host cell response and possibly dampen ISR, compared with other strains. Altogether, we assume that part of our observations is dependent on the biological context and may be relevant to conditions representative of the infection of the intestinal barrier by a clinical isolate of *Lm*, but obviously not of all possible occurrences of *Lm* intracellular invasion.

In addition to the gradual induction of ISR, our work highlights a strong translational repression of TOP-containing transcripts starting between 2 and 5 h p.i. The current model for the translational co-regulation of TOP mRNAs involves the direct binding of LARP1, a target of mTOR kinase activity, to TOP motifs^30^. When mTOR is inhibited, LARP1 becomes dephosphorylated and strongly binds TOP motifs, preventing mRNA translation initiation. However in our hands, we could only detect a consistent drop of mTOR kinase activity at 10 h p.i. (*Figure 3F*), which fails to explain the translational repression of TOP-containing transcripts observed at 5 h p.i. (*Figure 4*) unless these transcripts are more sensitive to mTOR inhibition than S6K1 phosphorylation is. The fact that we detected no phosphorylated form of LARP1 in non-infected LoVo cells (*Figure S7*) possibly indicates that other signalling pathways than the mTOR-phospho-LARP1 axis might be at play in the regulation of translation initiation of TOP-containing transcripts in our experimental context. It was also recently found that during exposure to sodium arsenite —a potent stress inducer that results in mTOR inhibition—, LARP1 was responsible for the recruitment of a fraction (10-15%) of TOP mRNAs to stress granules and P-bodies^31^. Whereas we observed that a small proportion of the TOP-containing mRNA *PABPC1* was associated with P-bodies (*Figure S8*), in our experimental conditions this proportion did not vary throughout the infection time-course, arguing against the docking of *PABPC1* to P-bodies being required for its translational repression. In the future, identifying and characterizing the molecular actors of the regulation of TOP-containing mRNAs in our experimental system could provide insights into the translational control of this class of transcripts in cells that are not metabolically hyperactive.

### Dynamics of gene expression response to *Listeria* infection

An important parameter addressed by our study is the timing of the host response to infection, and how different layers of gene expression control contribute to this timing. By studying a time-course of infection rather than a unique time point as had been done in most studies to date, and by quantifying not only the transcriptome but also the translatome, we reveal that in the first hours of infection most activation events are transcriptional, whereas most repression events are translational (*Figure 3A*). This rather binary effect is easily understandable by taking mRNA steady-state levels and turnover into account. Before infection, virtually no cytokine-encoding mRNA is present in cells; therefore their induction necessarily requires first transcription, and then translation. In contrast, mRNAs encoding components of the translation machinery are highly abundant. In addition, the intrinsic stability of human mRNAs is relatively high compared with the kinetics of an infection. Estimates of the median half-lives of mRNAs in mammalian cells are in the range of hours; among these, the half-lives of transcripts from housekeeping genes, especially those encoding translation factors and ribosomal proteins, are among the longest and often above 10 h^32,33^. As a consequence, downregulating the quantity of these transcripts by merely tuning down their transcription would take days. The most effective ways for a cell to rapidly stop the synthesis of proteins from this pathway would thus be either to massively degrade transcripts or to inhibit their translation. The second process — which we found to be prevalent between 2 and 5 h p.i. — is reversible; this might constitute an advantage during recovery from stress by allowing a rapid resumption of the translation of the previously-repressed genes. In the longer term, other types of regulation may take place; for instance, the downregulation of cytokine gene expression after 2 h p.i. likely relies both on the reduction of their transcription, and on the generally short half-life for this class of transcripts^33^. In addition to these broad rules drawn from the dominating patterns we observed, a number of individual transcripts are likely finely tuned by a combination of actions on transcription, decay and translation.

One level of regulation that was not addressed in our work was protein stability, which also should consistently contribute to the timing and effectiveness of the regulations we observed. Drastic alterations in the cell proteome and protein turnover in response to *Lm* infection or to treatment with LLO have previously been described^34,35^, and it would be interesting to integrate host proteomic data over an infection time course to assess how translation and proteome degradation contribute to reshaping the cell equipment. Within the time-course of our experiment, which was restricted to 10 h due to loss of viability afterwards, we have verified by immunoblot that the total amounts of PABPC1 were not affected (*Figure S9B*), probably due to the long half-lives of most core translation components^32,36^. In case the reduction of the amount of these proteins is playing a role by controlling the overall translation capacity of cells, it must thus be considered in a longer course of infection. In agreement with this, we did not monitor any consistent reduction in total protein synthesis activity within the 10 h we examined (*Figure 1*).

The translational repression of TOP-containing transcripts that we have observed is thus unlikely to impact the host cell proteome in a proportion that might affect infection during the first 10 hours. An alternative hypothesis would be that a reduction of the translation of TOP-containing mRNAs impacts infection outcome, rather than the reduction of the amounts of its products and of their functions. One could for instance imagine that, by pausing their anabolic metabolism and saving on the synthesis of abundant translation components, cells facing a bacterial challenge could reallocate part of their resources to antibacterial defences. Testing these hypotheses will deserve future investigations.

### Effects of the repression of *PABPC1* expression on bacterial or viral infections

Among the TOP-containing transcripts, *PABPC1* was the one most translationally repressed (*Figure 4*). The fact that pre-silencing its expression before infection reduced the intracellular multiplication rate of *Lm* suggests that the host cell can benefit from dampening its expression (*Figure 4L*). As discussed above for the bulk of TOP genes, this could either be due to a saving of resources by the host cell when avoiding the synthesis of a very abundant protein, or to a possible contribution of PABPC1 protein in the host-bacterial dialogue. Given the known function of PABPC1 as a regulator of mRNA translation and stability, its silencing is expected to have widespread effects on the host transcriptome and translatome, which could account for the decreased bacterial replication we observed when PABPC1 was knocked-down, apart from savings on host resources. This could include general effects due to the repression of overall translation, thereby impeding *de novo* protein synthesis and response to environmental cues^37^. The absence of PABPC1 could also impact genespecific regulations, as has been previously shown for developmental genes^38–40^, or genes containing A-rich motifs in their 5’-UTRs^41^. To the best of our knowledge, a specific regulatory function for PABPs in the response to a bacterial infection has not been addressed so far. And yet, a few pieces of evidence indicate that PABPs could contribute to the regulation of inflammation and innate immune responses. For instance, PABPC1 was among the top 5 differentially-expressed genes among patients with septic shock^42^. Blocking the ability of PABPs to bind poly(A) tails has been shown to reduce the sensitisation of mice to pain by dampening *de novo* protein synthesis and neurogenic inflammation in response to pro-inflammatory signals^37^. Direct interaction of Tristetraprolin/Zfp36 with PABPC1 was also found to mediate the translational repression of cytokine genes in PAMP-activated bone-marrow derived macrophages^43^. In response to viral infections, the product of an interferon-stimulated gene, RyDEN, was shown to restrict the replication of a variety of viruses by forming an inhibitory complex with PABPC1 and LARP1^44^. Both PABPC1 and LARP1 were positive regulators of Dengue virus (DENV) replication, reminiscent of our present findings for *Lm* intracellular multiplication. RyDEN was then hypothesized to interfere with DENV translation, which could not hold true in the case of a bacterial pathogen. A possible explanation to both phenotypes seen for *Lm* and DENV infections, and that would need testing, might be that PABPC1 inhibition would favour the expression of innate immune effectors that might help counteract infections.

## Conclusions

In response to *Lm* infection, the downregulation of host translation in LoVo epithelial cells affected a specific subset of transcripts, 5’-TOP-containing mRNAs, which encode components of the cell translation equipment. Among these, PABPC1 was the most repressed, and its down-regulation restricted intracellular replication. It appears reasonable to speculate that the coordinated translational repression of 5’-TOP-containing mRNAs would represent a host response to infection, rather than a bacterial strategy to subvert cell functions. The response to *Lm* infection thus contrasts with the translational repression observed during *Lp* infection^8^, which consistently affected the whole translatome of infected cells, and depended on the inhibitory function of at least four secreted *Lp* effectors on host translation elongation^7^. Whether effectors that interfere with host translation are also produced by *Lm* remains an open question.

## Material and methods

### Bacterial strains and culture conditions

The bacterial source strains used for this work were *Escherichia coli* NEB5α (New England BioLabs) for plasmid constructions, and *Listeria monocytogenes* LL195^17^ for all of the experiments involving *Lm*. All strains were grown at 37°C under shaking at 190 rpm in Luria Bertani (LB) medium for *E. coli*, in brain hear infusion (BHI) for *Lm*. Whenever required, media were supplemented with antibiotics for plasmid selection (chloramphenicol, 35 μg/ml for *E. coli;* 7 μg/ml for *Lm*).

For allelic replacement at the *hlyA* locus, the pMAD-Δ*hlyA* plasmid was created by amplifying two partially overlapping fragments by PCR: One thousand base pairs (bp) upstream (*plcA* gene) and downstream (*mpl* gene) of the *hlyA* open reading frame in the LL195 genome were amplified, respectively, with oligonucleotides oAL647-50 and oAL648-9 (*Table S6*). These fragments were inserted into the pMAD vector between the *Sal*I and *Bgl*II restriction sites by Gibson Assembly, using the NEBuilder HiFi DNA Assembly Cloning Kit (New England BioLabs). Allelic replacement of the *hlyA* open reading frame by this constructs, in the genome of *L. monocytogenes* strain LL195, was obtained as previously described^45,46^.

### Culture, infection and transfection of epithelial cells

Infections were performed in LoVo cells, an intestinal epithelial cell line originating from a colon adenocarcinoma (ATCC CCL-229). Cells were maintained in Ham’s F-12K (Kaighn’s) Medium (ThermoFischer Scientific Cat# 21127030), supplemented with 10% heat inactivated fetal bovine serum (PAN-Biotech) at 37°C and 5% CO_2_. Cells were passage 14 before seeding and were grown to 70-85% confluence prior to infection. The cell culture medium was changed every 24-h and kept for further use during the infection as “conditioned medium”. When needed, cells were transfected 48 h before infection with siRNAs against *PABPC1* (*Table S6*)^47,48^ or scrambled siRNAs using RNAiMAX in 24 well format as per manufacturer’s recommendations. Knockdown of targeted protein was confirmed by western blot. Control activation of the ISR pathway and inhibition of the mTOR pathway were obtained, respectively, by treating cells with 10 μg/ml of tunicamycin for 2 h or 20 nM of rapamycin for 1 h.

One colony of *Lm* was grown until they reached stationary phase (OD_600_ of 2 to 3) in BHI media at 37°C. Bacteria were washed with PBS, and added to a cell monolayer in the cell culture flasks (for Hi-seq, HPG incorporation or immunoprecipitation experiments) or in 24-well plate format (for gentamicin protection assay experiments) at a MOI of 30 to 40. The cell culture flasks or plates were centrifuged at 200 × *g* for 1 minute and then incubated at 37°C and 5% CO_2_ for 30 minutes. Cells were then washed with PBS containing 40 μg/ml gentamicin and conditioned media containing 25 μg/ml gentamicin was added. Infection was allowed to proceed until specific time points after which culture plates were snap-frozen in liquid nitrogen and stored at −80°C (for Hi-seq, HPG incorporation or immunoprecipitation experiments) or cells were washed in PBS and trypsinised for counting.

For gentamicin protection assays, cells were counted using a LUNA II automated cell counter and then centrifuged at 700g for 5 minutes. The cell pellet was re-suspended in water, incubated for 5 minutes, and then titrated through a 25G needle. Cell lysate was diluted in PBS and plated on BHI agar before overnight incubation at 37°C. Colony forming units (CFU) were counted and normalized to cell counts.

### Global translation using HPG incorporation and fluorescent labeling by copper catalyzed cycloaddition

L-Homopropargylglycine incorporation experiments were inspired from previous work exploring protein synthesis during HSV infection^49^. LoVo cells were grown to 70-85% confluence in 75 cm^2^ flasks and infected with *Lm* as described above. L-Homopropargylglycine (HPG, Jena Biosciences #CLK-016) was added at 2 mM final concentration 1 hour prior to each experiment end-point. Cells were snap frozen in liquid nitrogen and stored at −80°C. For copper-catalyzed alkyne-azide cycloaddition (click reaction), cells were lysed with click reaction compatible lysis buffer (100 mM HEPES, 150 mM NaCl, 1% Igepal CA-630) after which a click reaction was performed with Sulfo-Cy5-Azide (Jena Bioscience) at the following concentrations: 1 μg/μL protein, 100 μM azide sulfo Cy5 or azide-biotin, 1 mM Cu(II) sulfate, 5 mM Tris(3-hydroxypropyltriazolylmethyl)amine (THPTA), and 1 mM sodium ascorbate. Cu(II) sulfate was mixed with the THPTA prior to addition to the click reaction. Components were always added in the following order: azide-conjugate, Cu(II) sulfate-THPTA complex, and sodium ascorbate. The click reaction was allowed to proceed for one hour at room temperature. Samples were then methanol/chloroform precipitated and resuspended in Laemmli sample buffer (SB 1X). Samples were denatured at 95°C for 5 minutes and separated on a 12% SDS-PAGE gel. The gel was scanned with a Typhoon FLA 7000 biomolecular imager after which it was stained with colloidal Coomassie Brilliant blue G-250 as previously described^50^. Cycloheximide treated or no-treatment cells were used as negative controls.

Importantly, the cells were not starved of methionine prior to addition of HPG so that amino acid metabolism pathways remained unperturbed.

### Immunofluorescence and FISH on infected cells

LoVo cells were seeded in 24-well plates containing 12 mm diameter coverslips. Infection with bacteria expressing eGFP was performed as described above. At specified time-points, cells were fixed for 15 minutes with 4% paraformaldehyde/PBS, washed with PBS then stored at 4°C until further processing. Prior to staining, cells were permeabilized for 5 minutes at room temperature with 0.5% Triton X-100 in PBS. Cells were then blocked for 30 minutes in PBS buffer containing 2 % bovine serum albumin (BSA, Sigma) and incubated with Acti-Stain 670 fluorescent phalloidin (Cytoskeleton #PHDG1, 70 nM) and 4’,6-diamidino-2-phenylindole (DAPI, 100 ng/ml) for one hour. After three additional washes, cover glasses were mounted on microscope slides with Fluoromount mounting medium (Interchim). For PABPC1 FISH, a set of 48 Stellaris RNA FISH probes (Quasar^®^ 670 dye) against PABPC1 were designed using the Stellaris Probe Designer. PABCP1 mRNA FISH and immunofluorescent co-staining was done according to the Stellaris RNA FISH protocol. Antibodies and counter-staining were performed with DDX6 rabbit polyclonal primary antibody (Bethyl Laboratories #A300-460A), Cy3-goat-anti-rabbit secondary antibody (Jackson ImmuneResearch #111-165-144) both at a 1:500 dilution, and DAPI (100 ng/ml).

Preparations were observed with a Nikon Ti epifluorescence microscope (Nikon), connected to a digital CMOS camera (Orca Flash 4.0, Hamamatsu). Illumination was achieved using a SOLA-SE 365 source (Lumencor) and the following excitation/emission/dichroic filter sets (Semrock): DAPI, 377(50)/447(60)/FF409-Di03; Acti-Stain 670 or eGFP, 472(30)/520(35)/FF495-Di03. Images were acquired with Nikon apochromat 60x objective lenses (NA 1.4) and processed with the MicroManager and Fiji software. Each image is representative of the infected cell population.

### Immunoblotting

Total cell lysate was prepared by adding Laemmli sample buffer 1X supplemented with Pierce™ Universal Nuclease (Thermo Scientific™), phosphatase inhibitor cocktail and protease inhibitor cocktail directly to the cell monolayer. The monolayer was scrapped and the lysate was transferred to an Eppendorf tube after which samples were heated 95°C for 5 minutes and either stored at −80°C or used directly. Samples were migrated on a SDS-PAGE gel (12% acrylamide) and transferred to Amersham Hybond P 0.2 PVDF membranes using a Pierce G2 Fast Blotter (Thermo Scientific). Membranes were blocked in 5% w/v milk or BSA, TBS, 0.1% Tween^®^ 20 according to antibody manufacturer’s recommendations. Rabbit polyclonal antibodies (PABPC1, Atlas Antibodies #HPA045423; p70 S6 kinase α (H-160), Santa Cruz #sc-9027; LARP1, Bethyl Laboratories #A302-088A), rabbit monoclonal antibodies from Cell Signaling Technology (phospho-p70 S6 kinase (Thr389) clone 108D2 #9234; phospho-eIF2α (Ser51) clone D9G8 #3398) or mouse monoclonal antibodies (eIF2α clone D-3, Santa Cruz #sc-133132, β-Actin clone E4D9Z, Cell Signaling Technology #58169; Phosphoserine/threonine clone 22A, BD Transduction Laboratories # 612548) were added to the blocking solution at dilutions ranging from 1/500 to 1,5000 according to the manufacturer’s datasheets, and incubated overnight at 4°C. Membranes were incubated with corresponding secondary antibody (Bethyl Mouse or Rabbit IgG heavy and light chain antibodies coupled to HRP, # A120-101P and A90-116P) at a 1:50 000 dilution in the same buffer for 2 hours at room temperature. Signal was revealed using Pierce^®^ ECL Western Blotting Substrate on an ImageQuant™ LAS 4000 mini.

### Immunoprecipitation of LARP1

LoVo cells were grown to 80% confluence in 75 cm^2^ flasks and either left uninfected, or infected with *Lm* LL195 wild-type or Δ*hlyA* for 5 h as described above, or treated for 3 h with 100 nM of rapamycin^26^. Flasks were PBS-washed, snap frozen in liquid nitrogen and then stored at −80C° until further use. Total cell lysate was prepared by adding 150 μl of lysis buffer (40 mM HEPES pH 7.4, 120 mM NaCl, 1 mM EDTA, 1% Igepal CA-630) supplemented with 1X phosphatase inhibitor cocktail and protease inhibitor cocktail was added to each flask, and two flasks per condition were pooled. The obtained 300 μl of lysate were incubated 30 minutes on ice, disrupted by 10 passages through a 25G syringe needle, and clarified by centrifugation for 10 minutes at 12,000 × *g*, 4°C. 20 μl of each sample was kept as “input”, to which an equal volume of SB 2x was added. 2 μg of either LARP1 antibody, or control rabbit IgG were added to each sample, before overnight incubation at 4°C on a rotating wheel (10 rpm). 42 μg of Dynabeads-Protein G (ThermoFisher Scientific), pre-washed overnight in 10 volumes of IP buffer (40 mM HEPES pH 7.4, 120 mM NaCl, 1 mM EDTA, 1% Igepal CA-630), were added to each sample followed by incubation for 3 h at 4°C, 10 rpm. The flow-through fractions were discarder and beads were washed 3 times 2 minutes in 1 ml of IP buffer. Beads were finally resuspended in 25 μl of SB1X, denatured, separated on SDS-PAGE and probed by colloidal Coomassie staining and immunoblotting as described above.

### Haemolysis assay

Haemolysis titres of *Lm* LL195 and EGD-e strains were assessed as previously described^51^.

### RNA-seq and Ribo-seq sample preparation

LoVo cells were grown to 70-85% confluence in 75 cm^2^ flasks and infected with *Lm* LL195 as described above. Flasks were snap frozen in liquid nitrogen before infection (0 h) and at 2, 5, and 10 hours p.i., and then stored at −80C° until further use. Ribosome footprinting was done as per the protocol of Ingolia *et al*. Briefly, lysis buffer (20 mM Tris pH 7.4, 150 mM NaCl, 5 mM MgCl_2_, 1% TritonX, 1 mM DTT, 25 U/ml TurboDNase) was added to the frozen monolayer, which was then scrapped and transferred to an Eppendorf tube and processed. A portion of the lysate was taken, and acid phenol was used to extract total mRNA, which was stored at −80°C. Ribosome footprinting was performed on the same biological sample by adding RNAseI to the lysate at 2.5U/μl for 45 minutes at room temperature. The digestion was stopped by adding SuperaseIN (0.66U/μl) and ribosomes were purified by ultracentrifugation on a 1 M sucrose cushion. Ribosome protected fragments were extracted using acid phenol and used in sequencing library construction. Each time course was reproduced twice at a one week interval, thus producing biological triplicates.

### RNA-seq library construction

The IBENS Genomics Facility conducted the RNA-seq library construction. Isolated total cytoplasmic RNA integrity was verified using the RNA Pico method on the Agilent 2100 Bioanalyzer. High-quality RNA (RIN > 8) was used in library preparation for with the Illumina TruSeq stranded protocol (Illumina, San Diego, USA). Libraries were rRNA depleted using the Illumina Ribo Zero kit and sequenced as single read 75 base pair read length (SR75) on the NextSeq 550 system by the IBENS Genomics Facility.

### Ribo-seq library construction

Library construction was done using a protocol adapted from Huppertz *et al*. Briefly, RFPs were gel purified on polyacrylamide TBS-urea gels. The RFPs ends were then de-phosphorylated using T4 PNK and the 3’clip primer adaptor was ligated using T4 RNA ligase 2, truncated (New England Biolabs, #M0242). The ligated RFPs were gel purified and the RFPs were converted to cDNA using primers that contained barcodes and randomized nucleotides in order to remove PCR duplicates. The cDNA was then circularized using Circligase II (Lucigen, #CL4111K) and then linearized using BamHI. rRNA purification was performed as previously described^52^, with two modifications: (1) extra rRNA oligos were added to the biotinylated oligonucleotide cocktail, and (2) a second bead purification step was added. Purified RFP cDNA was amplified using Solexa primers and the libraries were sequenced as single read 75 base pair read length (SR75) on the NextSeq 550 system by the IBENS Genomics Facility.

The complete lists of oligonucleotides used for library constructions is supplied in *Table S6*.

### Read processing

RNA-seq reads that passed the Illumina Quality Filter (IQF) were aligned to rRNA (pre-rRNA 45S + rRNA 5S sequences from NCBI Nucleotide Database) using Bowtie2 (v2.3.2, option ‘-L 23’). Reads that were not mapped to rRNA were retained and aligned to the human genome (GRCh38) using the Spliced Transcripts Alignment to a Reference (STAR) software (v2.5.3a)^53^. Uniquely mapped reads were counted using featureCounts (v1.5.0)^54^.

For Ribo-seq reads, barcoded files were generated for each multiplexed fastq file. Reads that passed the IQF were then processed to remove PCR duplicates, which were identified by five random bases flanking the sample-specific barcodes. Reads that matched at these five random positions were classified as PCR duplicates and only the first hit was kept for further processing. Reads were trimmed (removing the 5’index and 3’adaptor) using cutadapt (v1.10) with option “-m/--minimum-length 25” to discard reads shorter that 25 nucleotides after adapter trimming. Trimmed reads were aligned to rRNA sequences as described above. The remaining reads were aligned to the human genome (GRCh38) using STAR (option ‘--*sjdbOverhang* 40’). Uniquely mapped reads were counted using featureCounts (v1.5.0).

All Hi-seq data discussed in this publication have been deposited in the European Nucleotide Archive^55^ and are accessible under accession number PRJEB26593 (https://www.ebi.ac.uk/ena/data/view/PRJEB26593).

### Data analysis and visualization

Library size normalized read alignments were visualized using the Integrative Genomics Viewer (IGV) from bedgraph files generated using samtools and bedtools. Post-mapping quality control and analysis of the distribution of reads by category of annotated genomic features was performed using ALFA^56^. Differential expression (RNA-seq) and differential translation (Ribo-seq) data were analyzed using DESeq2^57^. Translational efficiency was calculated and differential TE was analyzed using Riborex with a DESeq2 engine^58^. Functional enrichment analysis was conducted using over representation analysis of GO biological processes with the clusterProfiler R package^59^ on genes that had a false discovery *p*-value < 0.05 in the DESeq2 or Riborex analysis. Normalized Enrichment Score (NES) for TFBSs in the 500 bp region located upstream of the transcription start sites of RNA-seq DRGs were computed using RcisTarget^60^, which is R implementation of iRegulon^61^. Fuzzy clustering of TE values was performed using the Mfuzz package^62^. For the fuzzy clustering, TE values were recalculated by dividing TMM normalized (edgeR package) Ribo-seq counts by RNA-seq counts^63,64^. Only those genes that had an FDR p-value < 0.05 as computed by Riborex at any timepoint were included in order to decrease noise during clustering. ROAST (Rotation gene set tests for complex microarray experiments)^65^ was used for gene set testing of TOP-, uORF-, IRES-, or TISU-containing transcripts. The list of transcripts containing TOP motifs^66^, uORFs^67^ or IRES^68^ had been experimentally verified. In contrast, the list of transcripts containing a TISU motif with no mismatch was computed^69^. For correlative analysis between TE and 5’-UTR length, 5’-UTR sequences for protein coding transcripts were extracted from Ensembl using BioMart, and limited to those with MANE (matched annotation from NCBI and EMBL-EBI) Select annotation. Python 3.7 was used to parse the sequences into a data frame.

## Acknowledgements

We warmly thank Zhen Wang for sharing with us her optimized Ribo-seq protocol, and invaluable advice about how to best handle this technique. We are grateful to Caroline Peron Cane, Chloé Connan, Morgane Corre and José-Carlos Fernandez for their precious experimental assistance and eagerness to help solve technical issues, as well as Morgane Thomas-Chollier for helpful discussion regarding data analysis. Last, we thank the IBENS genomics platform for their careful sequencing of all samples, patient assistance and dedication, as well as the Computational Biology Centre for maintaining access to the servers, and Imaging facility for maintaining access to microscopy equipment.

## Author contributions

AL designed the project. VB and AL devised experiments and interpreted results. VB performed experiments with the help of SR for *Lm* genetics and FISH. FH, BN and VB analysed High-Seq data under supervision by AG and AL. AL and VB wrote the manuscript.

## Supplemental material

*Figures S1* to *S10* and *Table S6* are provided in this document. *Tables S1 to S5* being source data will be made available upon request.

**Figure S1.**
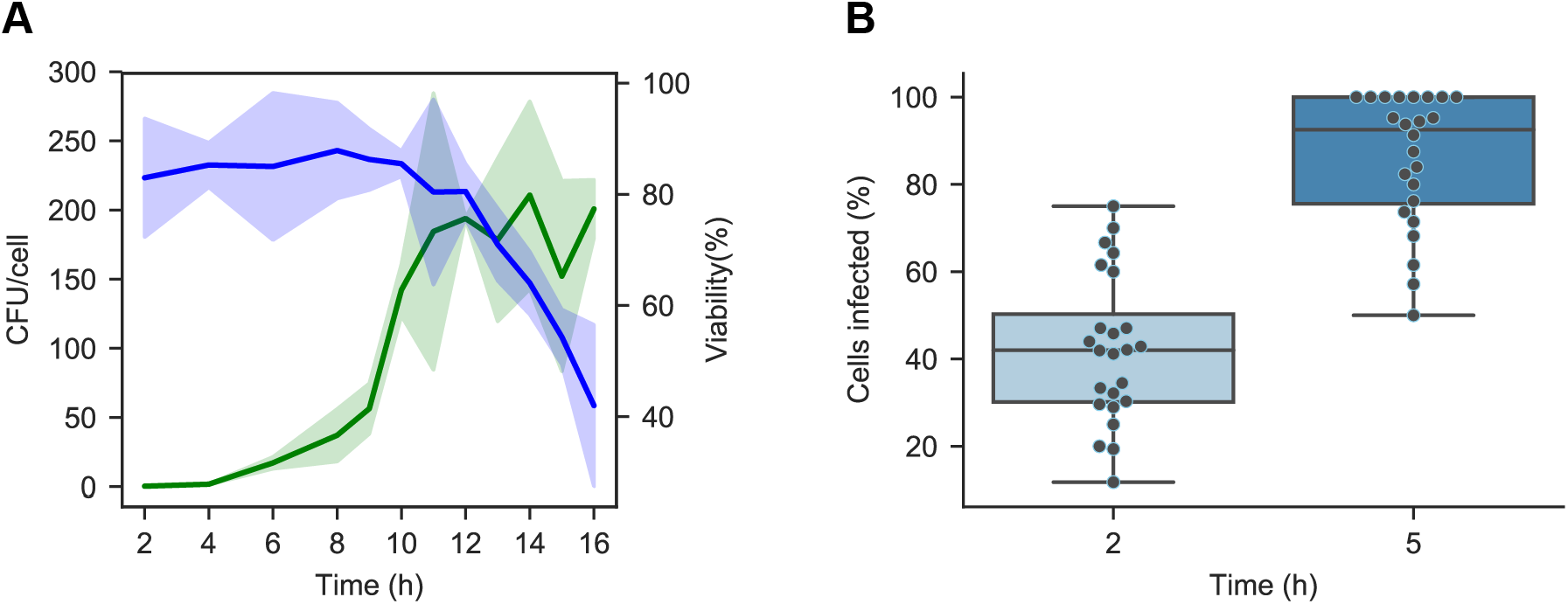
Assessment of infection kinetics and homogeneity. (**A**) The average number of bacteria per cell (CFU/cell, in green) and percentage of viable cells (Viability, in blue) were quantified over a 16-hour infection time course of LoVo cells, using an initial MOI of 30. CFUs were enumerated by plating serial dilutions on BHI-agar medium after cell lysis. Viability was assessed by live-dead staining using Trypan blue. Coloured bands in lighter shade indicate standard deviation. (**B**) Infection homogeneity was assessed by counting GFP-positive bacteria within each cell on microscopy slides, in 25 fields of vision per time-point. Cells were enumerated by counting DAPI-stained nuclei, and their contours were defined by revealing F-actin with Acti-Stain 670 fluorescent phalloidin. A cell was considered infected if it contained more than one bacterium.

**Figure S2.**
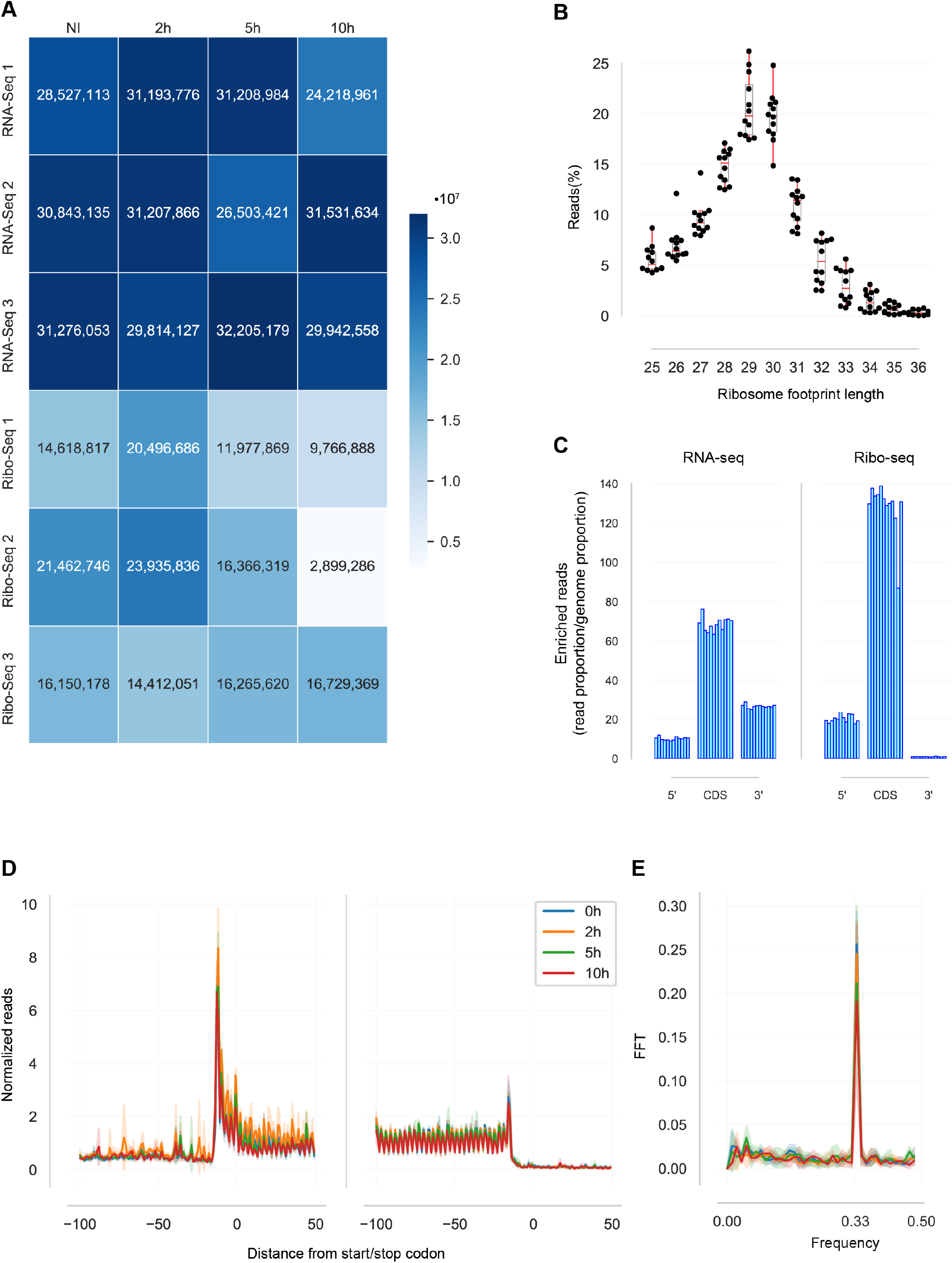
Quality controls of the RNA-seq and Ribo-seq data. (**A**) Number of uniquely mapped reads in each dataset reported in the present study. (**B**) Distribution of RFP read length. Each dot represents one of the Ribo-seq datasets. (**C**) Proportion of uniquely mapped reads matching to 5’-UTR, coding sequences (CDS) or 3’-UTRs in all transcripts across the RNA-seq (*left*) and Ribo-seq datasets (*right*) (**D**) Position of the 5’-end of sequenced RFPs relative to translation initiation sites (TIS) and stop codons in mRNAs. For each position within a 150-nucleotide region around the TIS or the stop codon, the counts of 5’-ends of RFP reads matching this position were summed across all transcripts in each dataset and normalized by the number of transcripts. Read 5’-ends were previously reported to match around 12-13 upstream of the ribosome A site, thus the signals resulting from ribosomes scanning CDSs typically start 12 nucleotides upstream of the TIS and terminate 12 nucleotides upstream of the stop codon^1^. (**E**) Codon periodicity of RFP reads in coding sequences. The signal decomposition by fast Fourier transform (FFT) of the average counts on open reading frames highlights a sharp peak at 0.33 frequency, indicative of a three-nucleotide periodicity in the RFP reads, consistent with ribosomes scanning the coding sequence during translation elongation. The colour code is as in (**D**).

---

**Figure S3.**
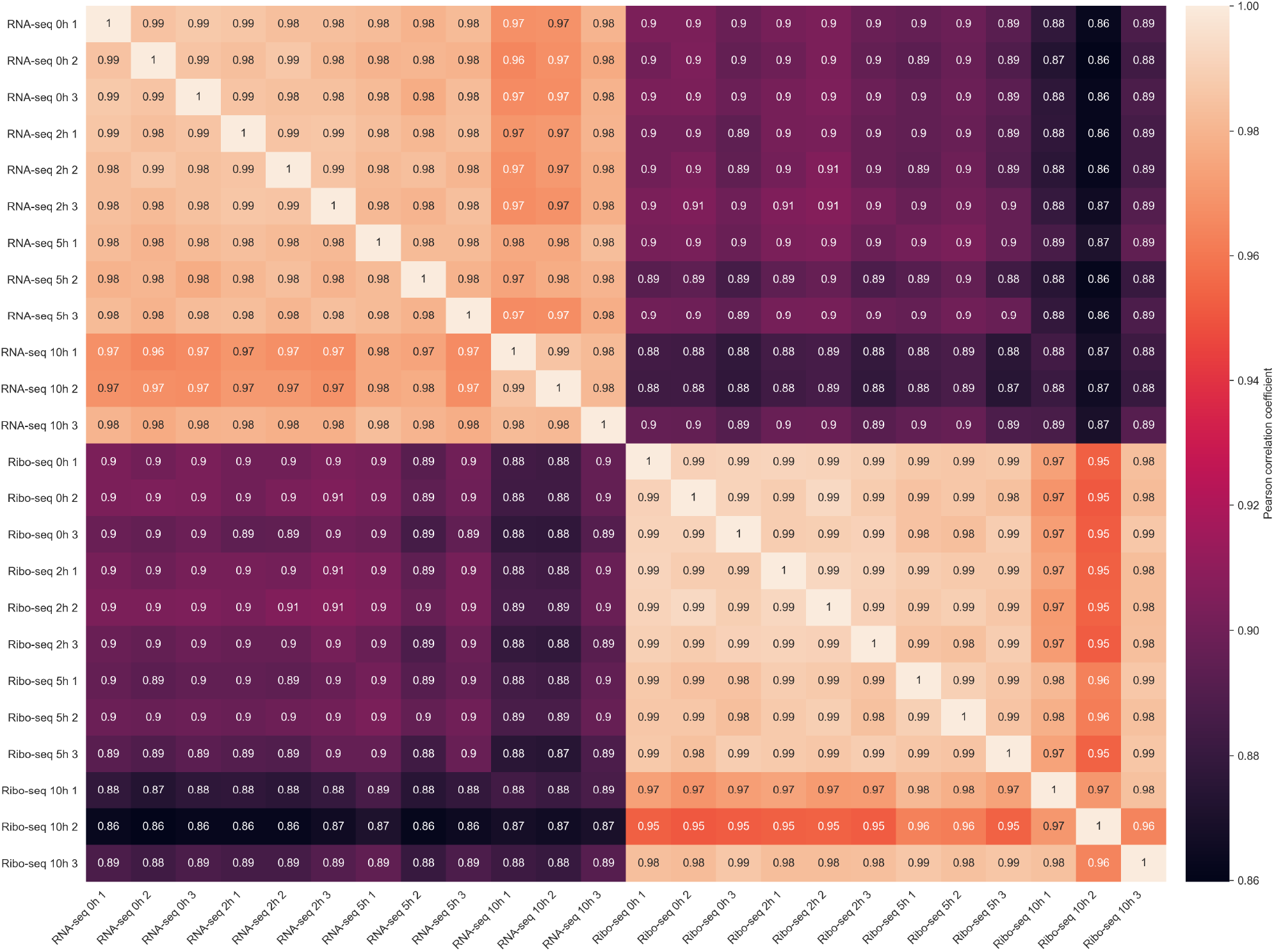
Correlative analysis of RNA-seq and Ribo-seq replicates. For each pair of RNA-seq or Ribo-seq samples, Pearson’s correlation coefficient of the Reads Per Kilobase of transcript, per Million mapped reads (RPKM) value for each gene was calculated. Lighter shades indicate higher correlation. Note that the Ribo-seq 10 h sample #2 stood out as an outlier, confirming our choice to exclude it for subsequent analysis. Other samples showed high correlation across RNA-seq experiments on the one hand, Ribo-seq experiment on the other hand.

**Figure S4.**
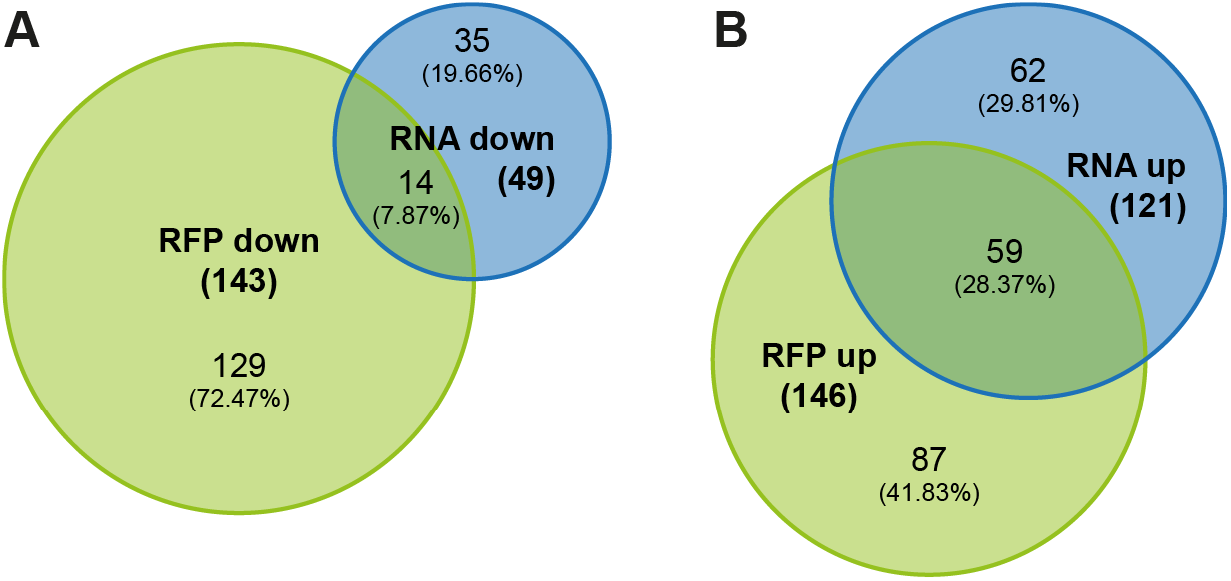
Overlap between differentially-regulated genes in the RNA-seq and Ribo-seq datasets between 2 and 5 h post-infection. The Venn diagrams highlight the number of differentially-regulated genes (DRGs with *p*_adj_ < 0.05 and FC > 1.5) in the RNA-seq (RNA, in blue) and Ribo-seq (RFP, in green) datasets, and their overlap. (**A**) down-regulated genes; (**B**) up-regulated genes. Percentages indicate the proportion out of the total number of genes on each panel. FC, fold change; *p*_adj_, adjusted *p*-value [DESeq false discovery rate (FDR)].

**Figure S5.**
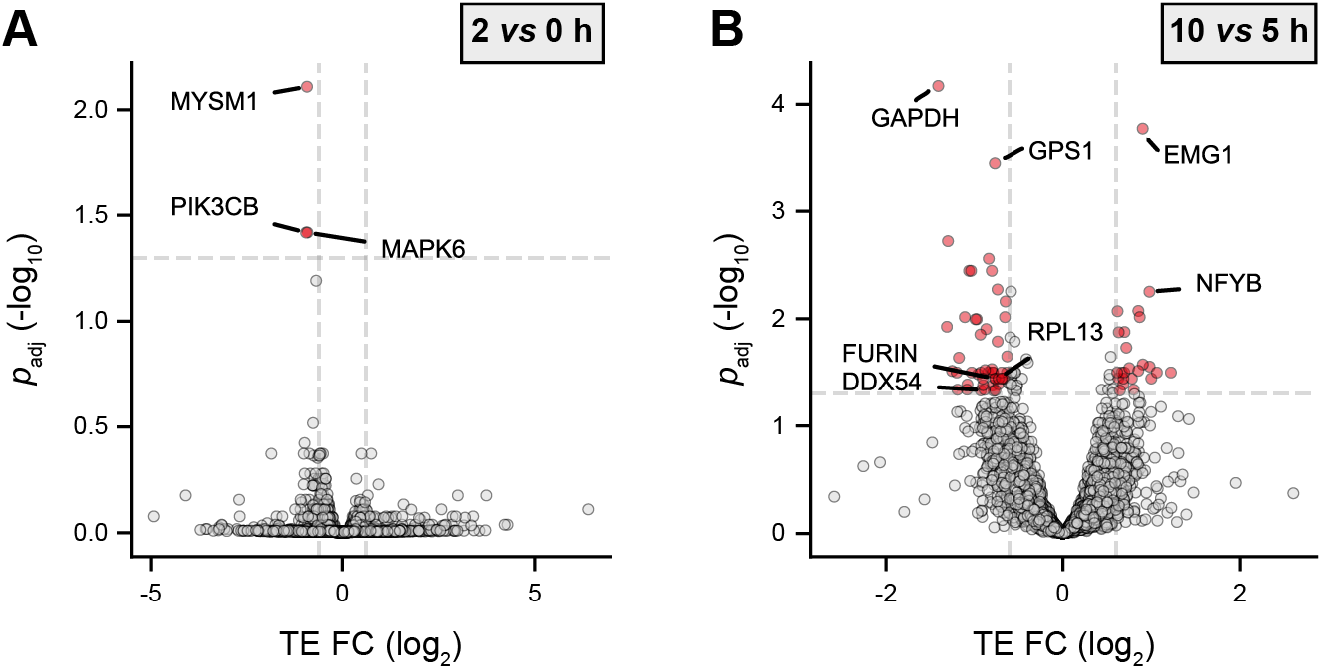
Variations in translation efficiencies during infection. The volcano plots highlight genes being significantly up- (*right*) or down- (*left*) regulated in translation efficiency (TE) at (**A**) 2 h p.i. compared to noninfected cells or (**B**) 10 h vs 5 h p.i. Data points coloured in red represent genes with an adjusted *p*-value below 0.05 (above dashed grey horizontal line; −log_10_ *p*_adj_ = 1.3) and a FC below or above 1.5 (vertical dashed grey lines; log_2_ FC = ± 0.58). Data from three independent replicates (except for RFPs at 10 h). FC, fold change; *p*_adj_, adjusted *p*-value.

**Figure S6.**
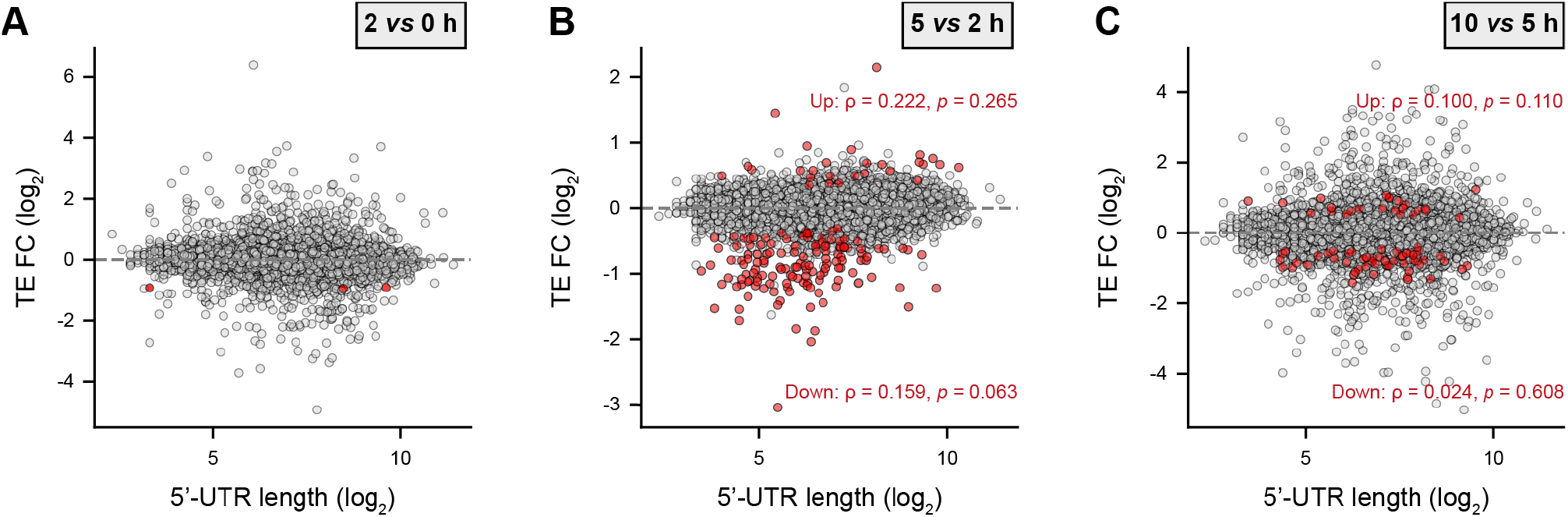
Correlation of changes in translational efficiency with 5’-UTR length. The log_2_ fold change (FC) of translational efficiencies (TE) between successive time-points of infection, calculated with Riborex, were plotted against the corresponding 5’-UTR length computed for each transcript with MANE Select annotation in Ensembl. Genes with an adjusted *p*-value on TE FC below 0.05 are highlighted in red. The Spearman’s rank correlation coefficients (p) and *p*-value for these significantly up- or down-regulated genes is indicated.

**Figure S7.**
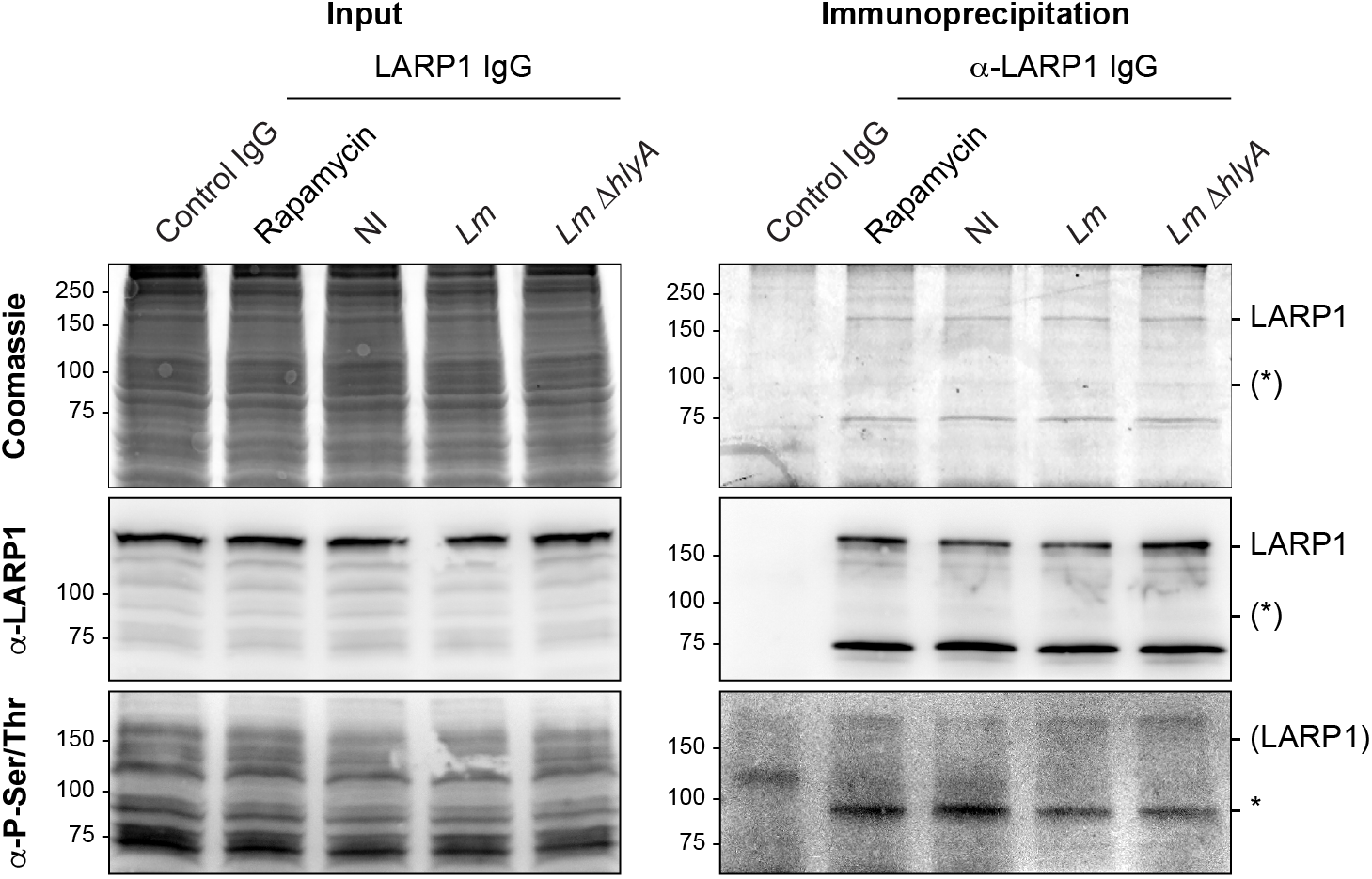
Phosphorylation status of LARP1 in LoVo cells. LARP1 was immunoprecipitated from LoVo cells that were infected or not for 5 h with *Lm* wild-type or with the *ΔhlyA* mutant strain, or treated for 3 h with 100 nM of rapamycin. Input and purified fractions were analysed by colloidal Coomassie staining (*top panel*) or by immunoblotting against LARP1 (*middle panel*) or using a total anti phospho-serine/threonine antibody (*bottom panel*). Note that whereas LARP1 appeared unphosphorylated in all samples, we observed that an unknown phoshoprotein of ~90 kDa (*, *lower panel*) co-immunoprecipitated with LARP1 in non-infected cells and in lower amounts (46% of non-infected) upon infection with either the wild-type or the *ΔhlyA* strains. The abundance and/or phosphorylation status of this protein in the co-immunoprecipitation was scarcely affected by rapamycin treatment (91% of non-infected), suggesting that it might be mTOR-independent. Identifying the nature of this LARP1-associated phosphoprotein and assessing whether it plays any role in the translation of TOP-containing transcripts in LoVo cells could represent a path for future mechanistic investigation of the translational regulations we observed in response to *Lm* infection.

**Figure S8.**
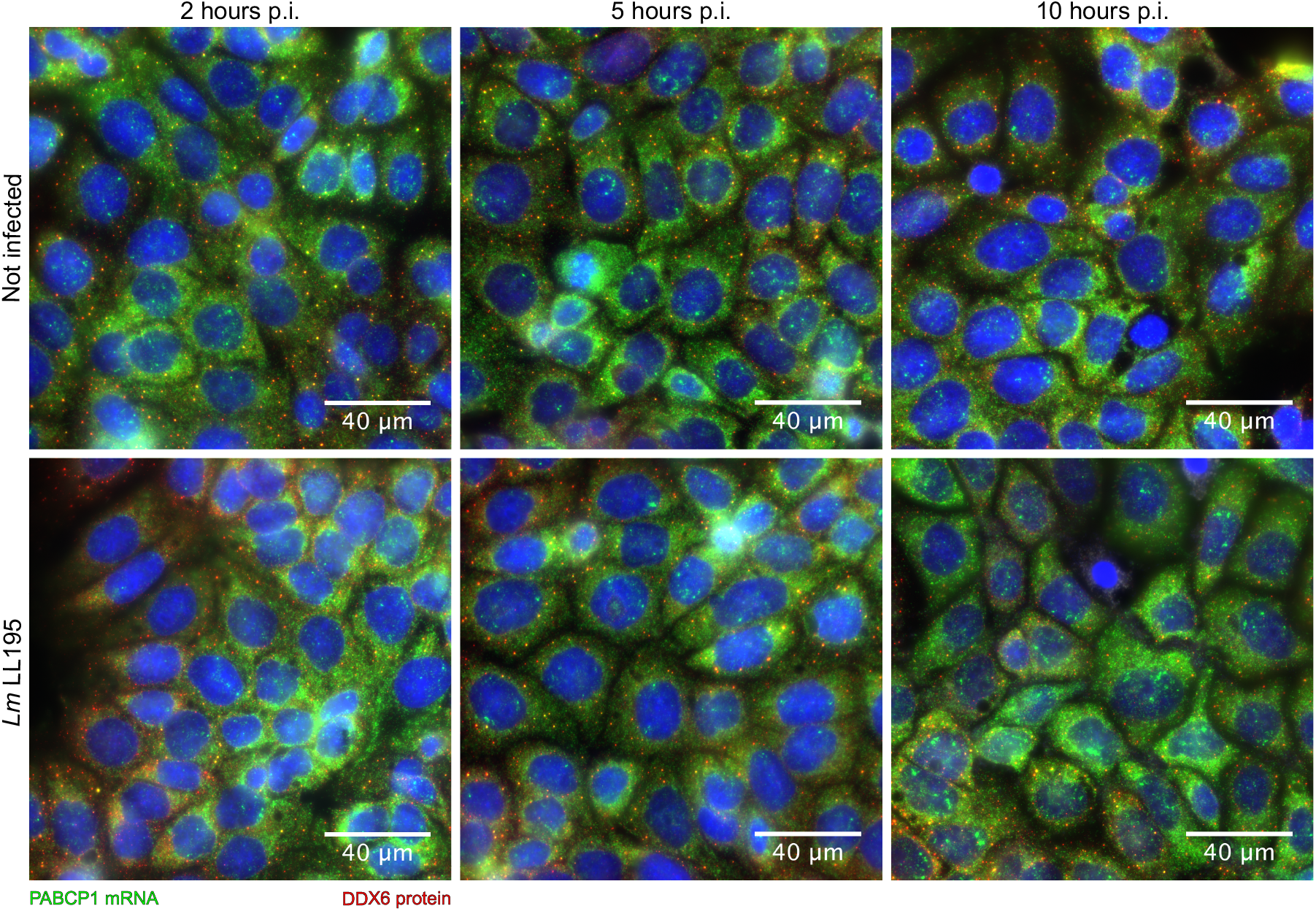
Localisation of *PABPC1* mRNA in LoVo cells infected or not with *Listeria monocytogenes*. LoVo cells were infected with *Lm* LL195 for 2, 5 or 10 h before fixation. The localisation of *PABPC1* mRNA was revealed by FISH (green), and that of P-bodies by immunofluorescence against DDX6 (red). Nuclei were stained with DAPI (blue).

**Figure S9.**
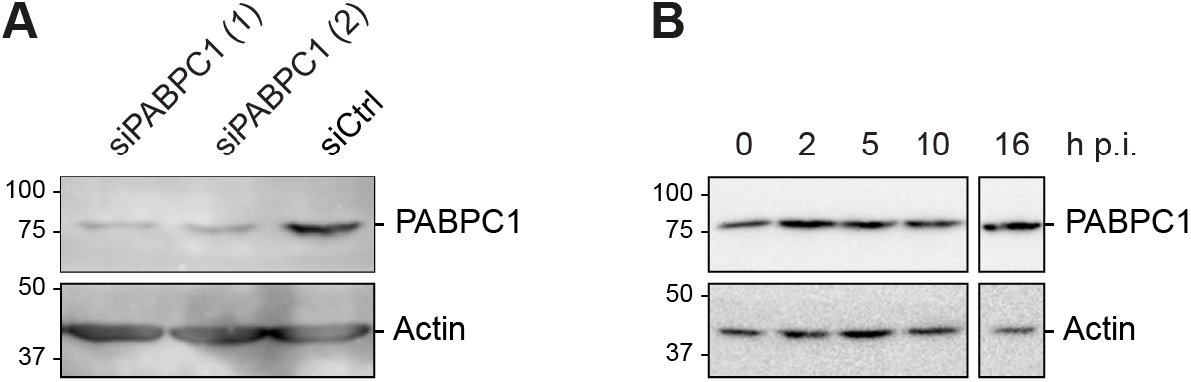
Assessment of PABPC1 protein levels upon transfection by siRNA or infection. PABPC1 and β-actin were revealed by immunoblotting. (**A**) *PABPC1* silencing by siRNAs. LoVo cells were transfected for 48 hours with siRNAs against *PABPC1* (siPABPC1 (1) and (2)) or with a scramble siRNA (siCtrl). (**B**) PABPC1 protein levels during infection. LoVo cells were infected for 0 to 10 h by *Lm* LL195 using a MOI of 30, or for 16 h using a MOI of 5.

**Figure S10.**
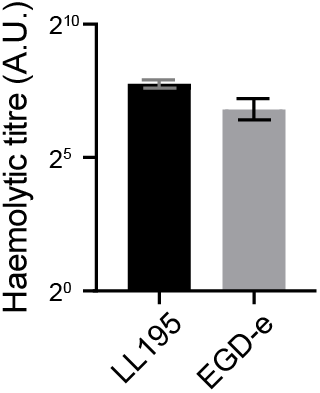
Haemolytic properties of *Listeria monocytogenes* strains LL195 and EGD-e. Data represent the means and standard deviations from three independent experiments.

**Table S1. Log2 FC and *p*_adj_ of variations in RNA, RFP and TE for each gene across infection time-points.**

**Table S2. Results of ORA of GO biological processes among DRGs in RNA-seq, Ribo-seq or TE at each time point of the infection.**

**Table S3. Top NES for TFBSs among DRGs in the transcritome datasets.**

**Table S4. Results of fuzzy clustering of TE variations over time, and ORA of GO biological processes on these clusters.**

**Table S5. Log2 TE FC for transcripts harbouring a TOP motif, a uORF, an IRES or a TISU in their 5’-UTR.**

**Table S1.**
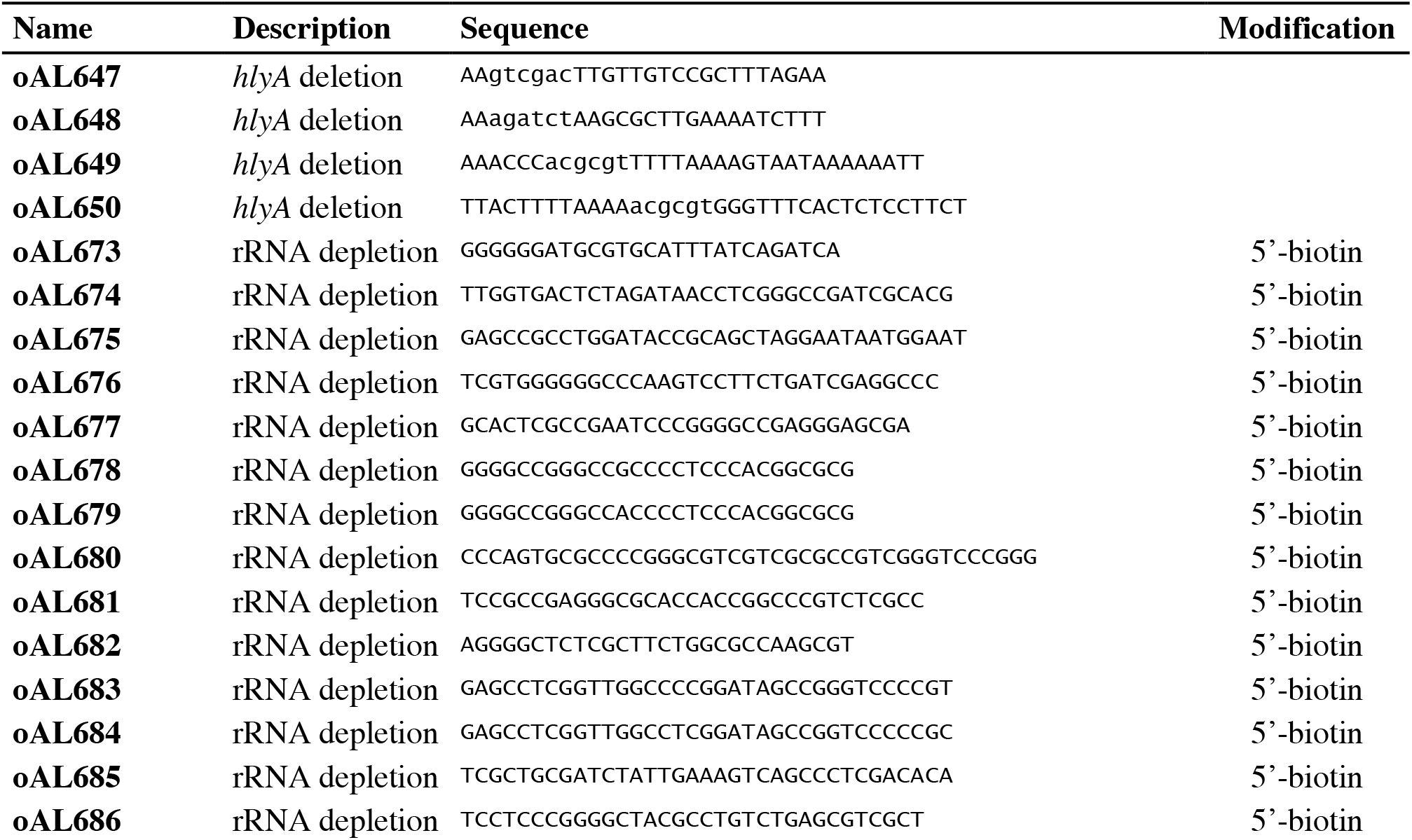
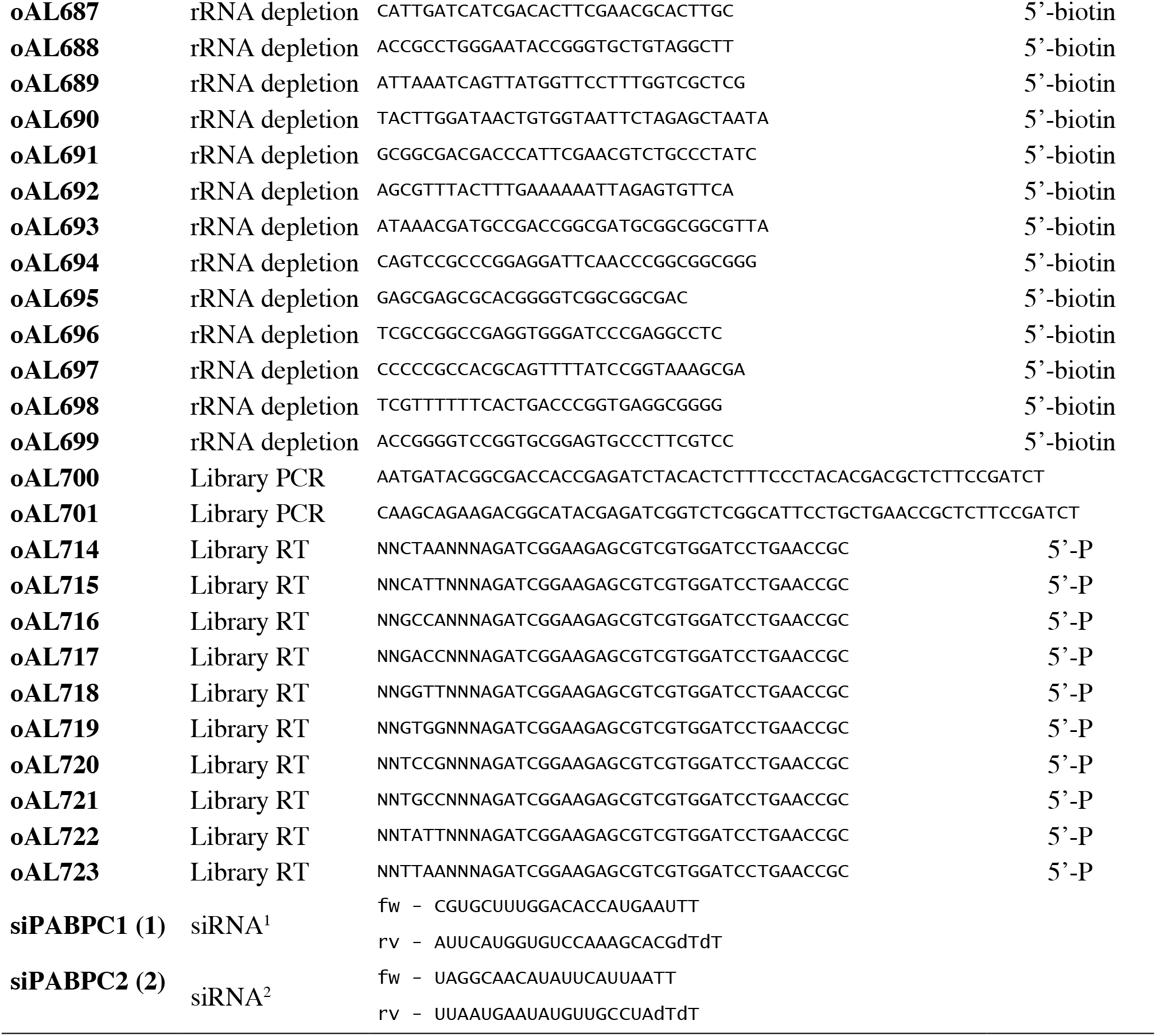
Oligonucleotides used in this study.

